# PKC-ε Regulates Vesicle Delivery and Focal Exocytosis for Efficient IgG-mediated Phagocytosis

**DOI:** 10.1101/2021.05.07.443102

**Authors:** Anna E. D’Amico, Alexander C. Wong, Cheryl M. Zajd, Ananya Murali, Michelle R. Lennartz

**Affiliations:** Albany Medical Center; Albany Medical College

## Abstract

PKC-ε is required for membrane addition during IgG-mediated phagocytosis; its role in this process is ill-defined. High resolution imaging revealed that PKC-ε exits the Golgi and enters phagosomes on vesicles that then fuse. TNF-α and PKC-α colocalize at the Golgi and on vesicles that enter the phagosome. Loss of PKC-ε and TNF-α delivery upon nocodazole treatment confirmed vesicular transport on microtubules. That TNF-α^+^ vesicles are not delivered in macrophages from PKC-ε null mice, or upon dissociation of the Golgi-associated pool of PKC-ε, implicates Golgi-tethered PKC-ε as a driver of Golgi-to-phagosome trafficking. Finally, we established that PKC-ε’s regulatory domain is sufficient for delivery of TNF-α^+^ vesicles to the phagosome. These studies reveal a novel role for PKC-ε in focal exocytosis: its regulatory domain drives Golgi-derived vesicles to the phagosome while catalytic activity is required for their fusion. This is one of the first examples of a PKC requirement for vesicular trafficking and describes a novel function for a PKC regulatory domain.

**Summary:** Golgi-tethered PKC-ε regulates vesicle trafficking along phagosomally-directed microtubules and vesicle fusion into forming phagosomes. Unexpectedly, the regulatory domain is sufficient for vesicle delivery.

**Graphical Abstract:** PKC-ε is required for Golgi-to-phagosome trafficking. Golgi-tethered PKC-ε^+^ vesicles carry TNF-α to IgG phagosomes. Dissociation of PKC-ε from the Golgi with PIK93 or expression of hSac1-K2A prevents vesicle delivery. The regulatory domain of PKC-ε (εRD) is sufficient for vesicle delivery, but not fusion. Created with BioRender.com

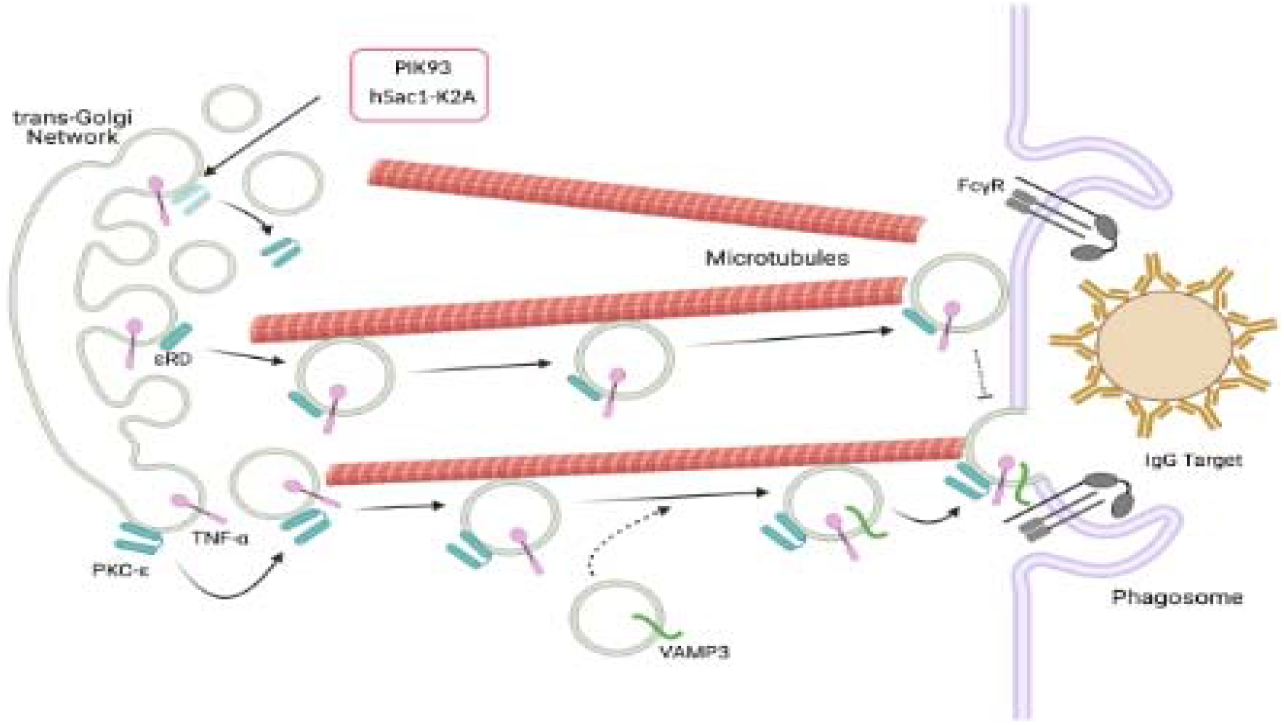

## Introduction/Background

Phagocytosis is the process by which innate immune cells, predominately neutrophils and macrophages, remove particulates from their environment. IgG-mediated phagocytosis is central to resolution of infection (Aderem, 2003; Niedergang and Chavrier, 2004). During phagocytosis, IgG-opsonized pathogens engage Fcγ receptors (FcγR), inducing receptor clustering and activation of downstream signaling networks that results in phagosome formation, closure, and pathogen clearance. Phagosome formation requires the addition of membrane for pseudopod extension and phagosome maturation (Botelho and Grinstein, 2011; Cannon and Swanson, 1992; Jaumouille and Grinstein, 2016; Lee et al., 2007). The membrane necessary for pseudopod extension is recruited from internal sources such as the endoplasmic reticulum and Golgi (Braun and Niedergang, 2006; Cannon and Swanson, 1992; Gerlach et al., 2020; Holevinsky and Nelson, 1998). Using patch clamping to quantify membrane addition, we reported that macrophages add ∼1/3 of their surface area from internal sources during spreading on IgG surfaces (ie, frustrated phagocytosis; (Wood et al., 2013). Work done by the Grinstein lab revealed vesicle associated membrane protein 3 (VAMP3) concentrates at the early phagosome, suggesting that targeted membrane delivery, or focal exocytosis, is involved in phagosome formation (Bajno et al., 2000).

Focal exocytosis is essential for various cellular processes including cell division, migration, and neurosecretion. The mechanisms underlying focal exocytosis have been well-studied in polarized cells such as neurons; however, little is known about this process in non-polarized cells. Phagocytosis, with its directed addition of membrane at phagosomes, provides a tractable model for studying focal exocytosis in non-polarized cells. Having demonstrated that Golgi-tethered Protein Kinase C-epsilon (PKC-ε) is required for membrane addition during FcγR-mediated phagocytosis (Hanes et al., 2017), this work was undertaken to determine the role of PKC-ε in the delivery and fusion of vesicles into the phagosome.

Protein kinase C’s are serine-threonine kinases grouped into three families (classical, novel, and atypical) based on their activators. They share a common domain structure, a unique regulatory domain tethered to a homologous kinase domain by a flexible hinge. Within the regulatory domain is a pseudosubstrate region (PS) that binds in the active site, maintaining the enzyme in an inactive conformation, a C1 region that binds diacylglycerol (DAG), and for Ca^2+^ dependent isoforms, a C2 Ca^2+^-binding region (Newton, 1997). Upon cell activation, they translocate to membranes, binding their activators, resulting in a conformational change that releases the pseudosubstrate, exposing the active site for substrate phosphorylation (Newton, 1997). In resting cells, PKC-ε is predominately cytosolic (Larsen et al., 2000).

We identified PKC-ε, one of the novel isoforms, as a critical player in membrane mobilization during phagocytosis (Cheeseman et al., 2006; Newton, 1997). PKC-ε is unique in its substrate specificity as demonstrated in in vitro binding and chimeric studies (Pears et al., 1991; Schaap et al., 1989). We reported that chimeras of the PKC-ε regulatory domain linked to the PKC-δ catalytic domain concentrate at phagosomes but fail to promote phagocytosis, consistent with the unique substrate specificity of PKC-ε (Wood et al., 2013). Additionally, we reported that PKC-ε is tethered to the Golgi through binding of its pseudosubstrate domain to PI4P (Hanes et al., 2017). Dissociation of this Golgi-associated PKC-ε pool using the PI4-kinase inhibitor PIK93, or expression of the Golgi-targeted PI4-phosphatase hSac1-K2A, abrogates PKC-ε concentration at the phagosome and slows the rate of phagocytosis, suggesting that Golgi-associated PKC-ε is involved in membrane mobilization. Patch-clamping of wild type and PKC-ε null macrophages during frustrated phagocytosis revealed that virtually all of the membrane added in response to FcγR ligation is dependent on PKC-ε (Wood et al., 2013)That dissociation of the Golgi-tethered pool of PKC-ε with PIK93 blocked this membrane addition established a PKC-ε dependent Golgi-to-phagosome vesicular trafficking highway (Hanes et al., 2017). These data allude to a novel mechanism of PKC translocation in which PKC-ε translocates to its site of action from the Golgi rather than the cytosol.

In this study, we investigate the mechanism underlying the PKC-ε trafficking from the Golgi for focal exocytosis during FcγR-mediated phagocytosis. Our results demonstrate that PKC-ε traffics on microtubules associated vesicles from the Golgi to the forming phagosome. Furthermore, PKC-ε, more specifically the regulatory domain of PKC-ε, is *required* for Golgi-to-phagosome vesicle movement.

## Results

### PKC-ε Enters the Phagosome on Vesicles

Our previous work identified PKC-ε as a critical component of the signaling network for IgG-mediated phagocytosis. PKC-ε concentrates at the forming phagosome (Larsen et al., 2002) and PKC-ε null macrophages have significantly smaller pseudopods and slower phagocytosis (Hanes et al., 2017; Wood et al., 2013). More recently, we established that a pool of PKC-ε is tethered to the trans-Golgi network (TGN) through binding of its pseudosubstrate domain to PI4P (Hanes et al., 2017). PKC-ε can be dissociated from the TGN with the PI4 kinase inhibitor PIK93 or by expressing the Golgi-directed PI4 phosphatase, hSac1-K2A (Mayinger, 2009). PKC-ε fails to localize at bound targets and phagocytosis is significantly slower in macrophages lacking TGN-associated PKC-ε. Finally, using patch-clamping to quantitate membrane capacitance in macrophages treated with PIK93, we established that the TGN pool of PKC-ε is required for membrane addition in response to FcγR ligation. *How* Golgi-associated PKC-ε regulates membrane addition at the phagosome remains an open question. However, these data are consistent with a model in which PKC-ε vesicles fuse to expand the membrane for target internalization.

Live imaging of PKC-ε-GFP expressing macrophages during IgG-mediated phagocytosis revealed PKC-ε puncta exiting the Golgi (Figure 1A and Video 1). That Golgi-associated PKC-ε concentrates at the phagosome (Hanes et al., 2017), coupled with visualization of PKC-ε export from the Golgi as puncta, is evidence that the PKC-ε localizing at the phagosome originates at the Golgi, and implicates vesicular trafficking as its mode of transport. Total Internal Reflection Fluorescence Microscopy (TIRFM) during frustrated phagocytosis (i.e, spreading on IgG surfaces) can distinguish vesicle appearance (puncta) at the membrane vs cytosolic translocation (gradual homogenous increase) (Figure 1B and Videos 2-4). The pattern of PKC-ε-GFP appearance at the membrane was compared to that of VAMP3-GFP (a transmembrane protein delivered to phagosomes on vesicles) and Akt-PH-GFP (PH domain of Akt), a lipid reporter that translocates from the cytosol to bind PIP_3_ at the phagosome (Bajno et al., 2000; Niedergang and Chavrier, 2004; Yeo et al., 2015). Their patterns of entry were distinct. Akt-PH-GFP fluorescence at the membrane increased homogenously and was lost over time (Figure 1B, top panel) while VAMP3-GFP appeared as distinct puncta (Figure 1B, middle panel). PKC-ε-GFP also appeared as puncta (Figure 1B, bottom panel), indicating that PKC-ε enters the phagosome on vesicles. Because PKC-ε and VAMP3 present with a similar (punctate) pattern, we asked whether PKC-ε and VAMP3 colocalize. BMDM expressing either VAMP3-GFP or Akt-PH-GFP were spread on IgG surfaces, fixed, and stained for endogenous PKC-ε (Figure 1C). Pearson’s correlation coefficient revealed significantly higher colocalization between PKC-ε and VAMP3-GFP compared to PKC-ε and Akt-PH-GFP (Figure 1C, graph). Colocalization of PKC-ε with a transmembrane protein confirms its presence on vesicles. Additionally, the punctate patten of *endogenous* PKC-ε matched that of PKC-ε-GFP, validating that PKC-ε-GFP is a valid readout of its endogenous counterpart.

**Figure 1:**
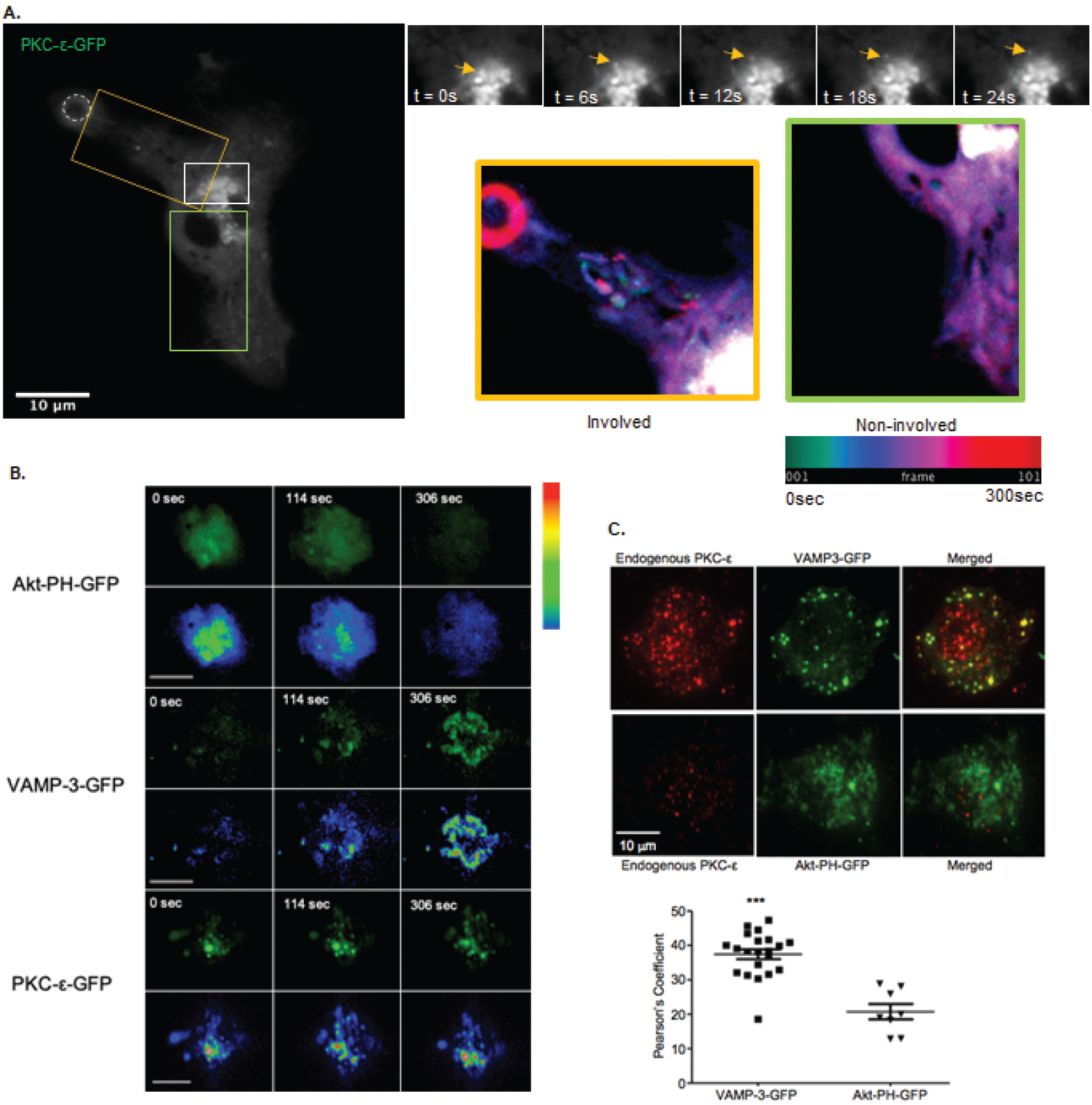
PKC-ε Traffics on and Enters the Phagosome on Vesicles. (A) BMDM, transduced to express PKC*-ε-*GFP, were presented with 5μm IgG-opsonized particles and phagocytosis was followed with time. Images were captured at 3 sec intervals over 10 min by spinning-disc confocal microscopy. Puncta can be seen exiting the Golgi (arrows). Time colored puncta tracks are seen directed towards the phagosome but are absent from non-phagocytic regions of the cell. Temporal-color coding was applied to images post-acquisition using FIJI Representative of 15 cells collected from 5 independent experiments. (B) BMDM, expressing Akt-PH-GFP, VAMP3-GFP, or PKC-ε -GFP were followed with time as they attached and spread on lgG surfaces, Images were captured by TIRFM every 8 seconds for 10 min. A pseudocolored fire LUT was applied to images for optimal visualize of fluorescence intensity (bottom panels. Representative frames from time-lapse imaging (Videos 2-4) are presented: 0 time is set as the first frame in which fluorescence appears. Representative of 4 experiments. PKC-ε presents as puncta, more similar to (transmembrane VAMP3 than (cytosolic) Akt-PH. (C) BMDM expressing GFP conjugated VAMP3 (upper) or Akt-PH (lower) were allowed to bind to IgG surfaces (5 min, 37°C) fixed, stained for endogenous PKC-ε, and imaged by TIRF. The colocalization between PKC-ε and VAMP3-GFP or Akt-PH-GFP, presented as the Pearson’s correlation coefficient, revealed low association with Akt-PH but significantly higher colocalization between PKC-ε and VAMP3 (graph). Data are presented as mean ± SEM with each symbol representing one cell with the aggregate data from 3 independent trials. Significance tested by Students’ t-test: ***p<0.005 for VAMPS compared to Akt. Scale bars = 10 μm

If the PKC-ε that appears at the phagosome originates from the TGN, then depletion of the Golgi pool should reduce PKC-ε puncta at the membrane. PIK93, the PI4K II inhibitor that depletes Golgi-tethered PKC-ε (Hanes et al., 2017) reduced the number of PKC-ε-GFP puncta at the phagosome 6-fold compared with untreated controls (Figure 2). The effect was reversable as puncta delivery recovered upon PIK93 washout. The PIK93 effect was not due to the disruption of the Golgi integrity, as it did not alter the pattern of GM130 (cis-marker) nor Golgin-245 (TGN marker) staining (Supplemental Figure 1). Notably, the GFP signal at the membrane in PIK93 treated cells showed very few puncta and no substantive increase in membrane intensity, supporting the conclusion that the puncta at the membrane are derived from the Golgi and there is little to no contribution from the cytosolic PKC-ε pool.

**Figure 2.**
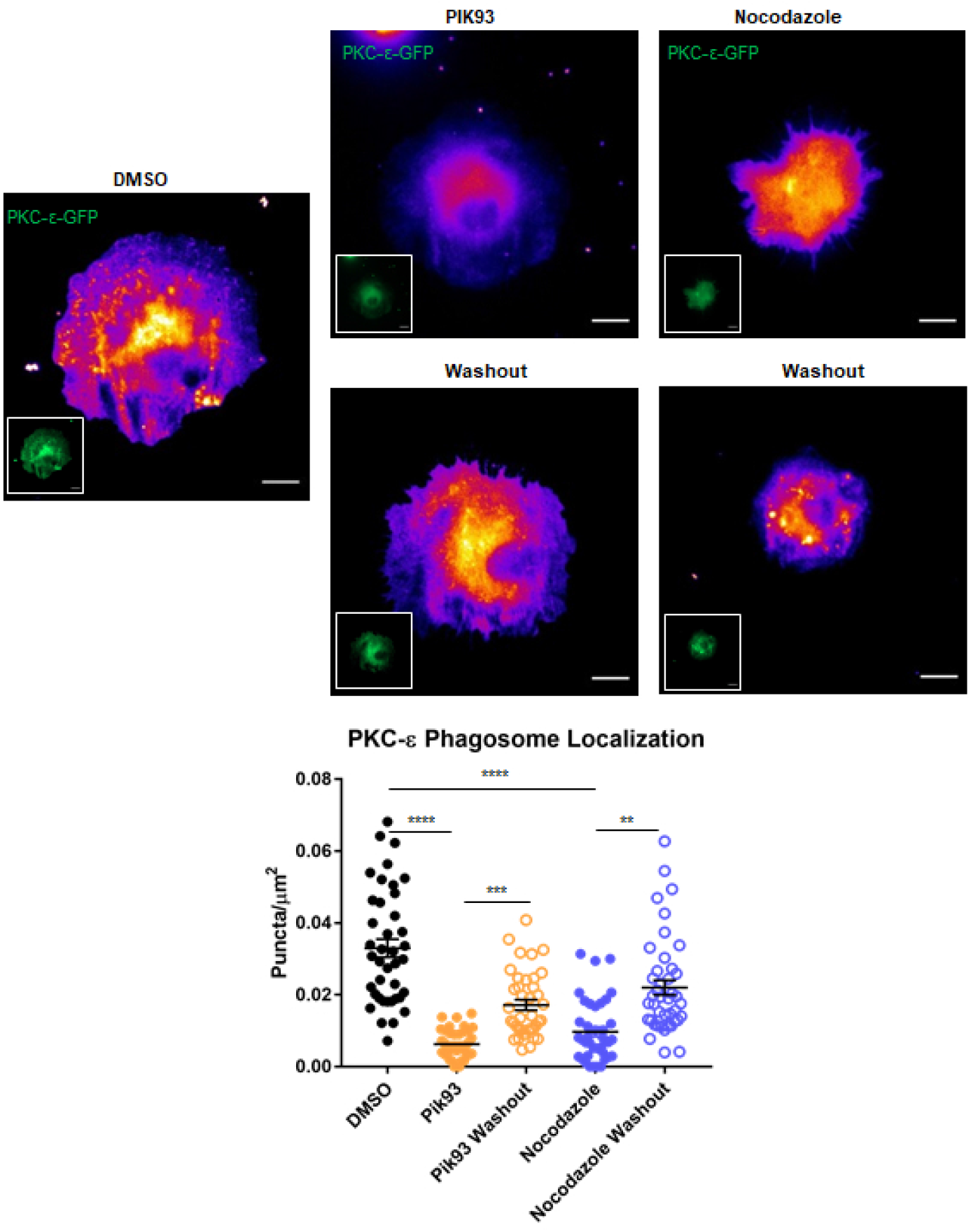
PKC-ε delivery is reversibly inhibited by nocodazole or PIK93. BMDM expressing PKC-ε -GFP were treated with DMSO (Solvent control), PIK93 (100nM), or nocodazole (10 μM) for 30 min then spread on IgG surface for 30 min and fixed. For washouts, the inhibitor-containing media was replaced with HBSS ^+ +^ for 5 min prior to fixation. Fire LUT was applied for optimal visualization of puncta, inserts represent GFP expression. Samples were imaged by TIRFM and the number of puncta was quantitated by FIJI software. Data are presented as number of puncta/cell area to account for differences in cell spreading. Data are presented as mean ± SEM of 3 independent experiments (total of 39-45 cells per condition), each data point represents a cell. Statistical significance was determined by one-way ANOVA with Bonferroni post-test. ****p<0.0001, ***p<0.001, **p<0.01. Scale bar = 10 μm.

If PKC-ε transits from the TGN to the phagosome on vesicles, microtubules likely serve as the highway for their transport. Consistent with this, we reported that the microtubule disrupting agent nocodazole prevents membrane addition in response to FcγR ligation (Hanes et al., 2017). But does it block PKC-ε vesicle delivery? Indeed, fewer PKC-ε^+^ vesicles were delivered in nocodazole-treated BMDM, an effect that was rescued upon drug washout (Figure 2, right panel). Together, these data provide the first evidence that a PKC, any PKC, translocates to its site of action on a vesicle. Specifically, the results show that the PKC-ε that concentrates at phagosome originates in the Golgi and transits on vesicles in microtubule-dependent manner.

### PKC-ε Vesicles Fuse into the Forming Phagosome

While the above experiments demonstrate that vesicles dock at the membrane, they do not address their fusion. If the Golgi pool of PKC-ε is required for membrane addition, (Hanes et al., 2017) and if PKC-ε traffics to the phagosome on vesicles (above), we would predict that the PKC-ε^+^ vesicles that approach the membrane actually fuse. When a fluorescent vesicle approaches the membrane, its intensity by TIRMF increases. If the vesicle fuses, the peak intensity will decrease as the fluorescent probe diffuses into the plane of the membrane, resulting in an increase in mean intensity. Thus, vesicle fusion is defined as a rise (approach) and fall (fusion) in maximum fluorescence intensity combined with an increase in overall fluorescence intensity (Jaiswal et al., 2009; Schmoranzer et al., 2000). Thus, we tracked PKC-ε-GFP as BMDM attached and spread on IgG surfaces and used post-acquisition analytics of puncta region of interest (ROI) to determine the maximum (peak) and overall (mean) fluorescence intensity of vesicles with time (Figure 3 and Video 5). Over time, the peak fluorescent intensity increased. The subsequent decrease in peak intensity was accompanied by an increase in overall fluorescence in the ROI (Figure 3B and C, left panel and Supplemental Figure 2), fulfilling the established criteria for vesicle fusion (Jaiswal et al., 2009; Schmoranzer et al., 2000). Analysis of an equivalent ROI lacking vesicles in the same cell served as an internal control and showed no change in peak or mean fluorescence intensity (Figure 3C, right panel and Supplemental Figure 2). These data demonstrate that PKC-ε^+^ vesicles fuse into the forming phagosome.

**Figure 3:**
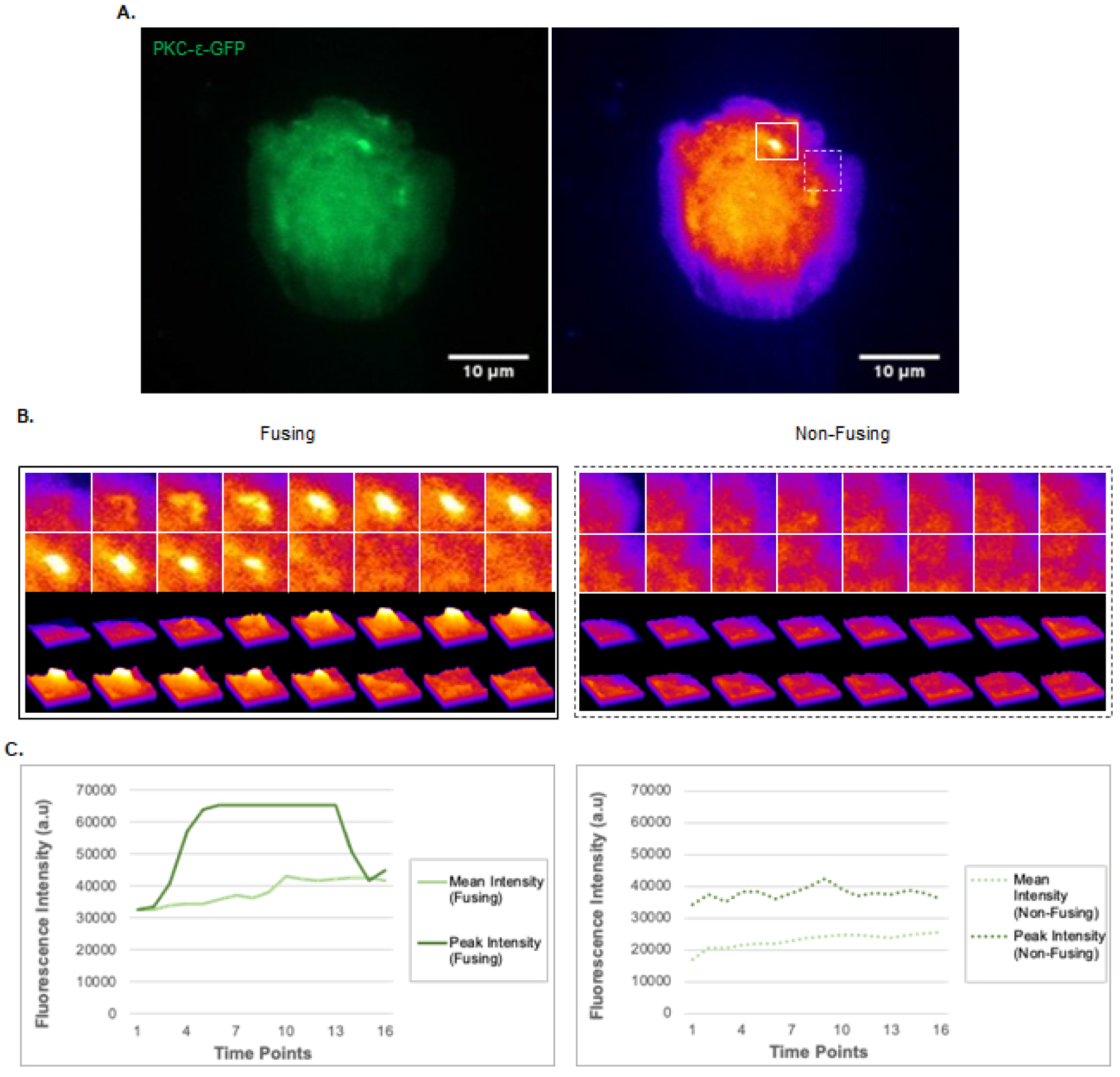
PKC-ε ^+^ Vesicles Fuse into the Phagosome. BMDM expressing PKC-*ε* -GFP were followed with time as they attached and spread on IgG surfaces, Images were captured by TIRFM every 8 seconds for 10 min. (A) Single timepoint image from the video (Video 5) is shown with an ROI containing a vesicle (solid box) and an equivalent area lacking vesicles (dotted box). (B) Pseudo-coloring of the ROIs was applied for optimal visualization of fluorescence intensity of fusing and non-fusing (no vesicle) ROls. Upper panels of show intensity as a LUT while lower panels use a 3D visualization of intensity. (C) Quantitation of mean and peak fluorescence intensity was done using FIJI software. As a vesicle approaches the surface, the intensity increases (peak intensity) and decreases upon fusion, spreading the fluorescence within the plan of the membrane increase the mean intensity in the ROI). Representative of 22 fusion events from 2 independent experiments. Scale bar = 10μm. Supplemental Figure 2 shows other fusion events within the same cell.

### PKC-ε Vesicles Align Along Microtubules

The fact that nocodazole prevents membrane addition (Hanes et al., 2017) and delivery of PKC-ε vesicles to the phagosome (Figure 2) suggests that PKC-ε^+^ vesicles travel on microtubules. Thus, we determined the extent to which PKC-ε aligns with microtubules in BMDM in response to FcγR ligation. BMDM were subjected to frustrated phagocytosis, fixed, stained for endogenous PKC-ε and α-tubulin, and imaged by Airyscan high resolution microscopy (Figure 4A and Supplemental 3A). Two different methods of alignment analysis were used. First, we traced either along α-tubulin tracks (Fig 4A and B, solid lines 1-3, and Supplemental Fig 3A) or across tracks (Fig 4A and B, dotted lines a-c), to look for co-localization of PKC-ε puncta with α-tubulin. If PKC-ε was associated with MT, we reasoned that we would see co-localization of (red) PKC-ε with (green) α-tubulin. Line plots of PKC-ε and α-tubulin fluorescence were generated on a single slice from a z-stack image using FIJI software. Regardless of how the lines were drawn, the majority of PKC-ε fluorescence co-localized with α-tubulin (Figure 4B and Supplemental 3A), suggesting PKC-ε vesicles associate with microtubules. Secondly, whole-cell 3D reconstructed images were generated and, using Imaris software, spots and surfaces were constructed for PKC-ε and α-tubulin, respectively. Analysis revealed the ∼52% of PKC-ε puncta were aligned along microtubules (color-coded pink spots in Figure 4C and Supplemental 3B), consistent with PKC-ε puncta trafficking on microtubules.

**Figure 4:**
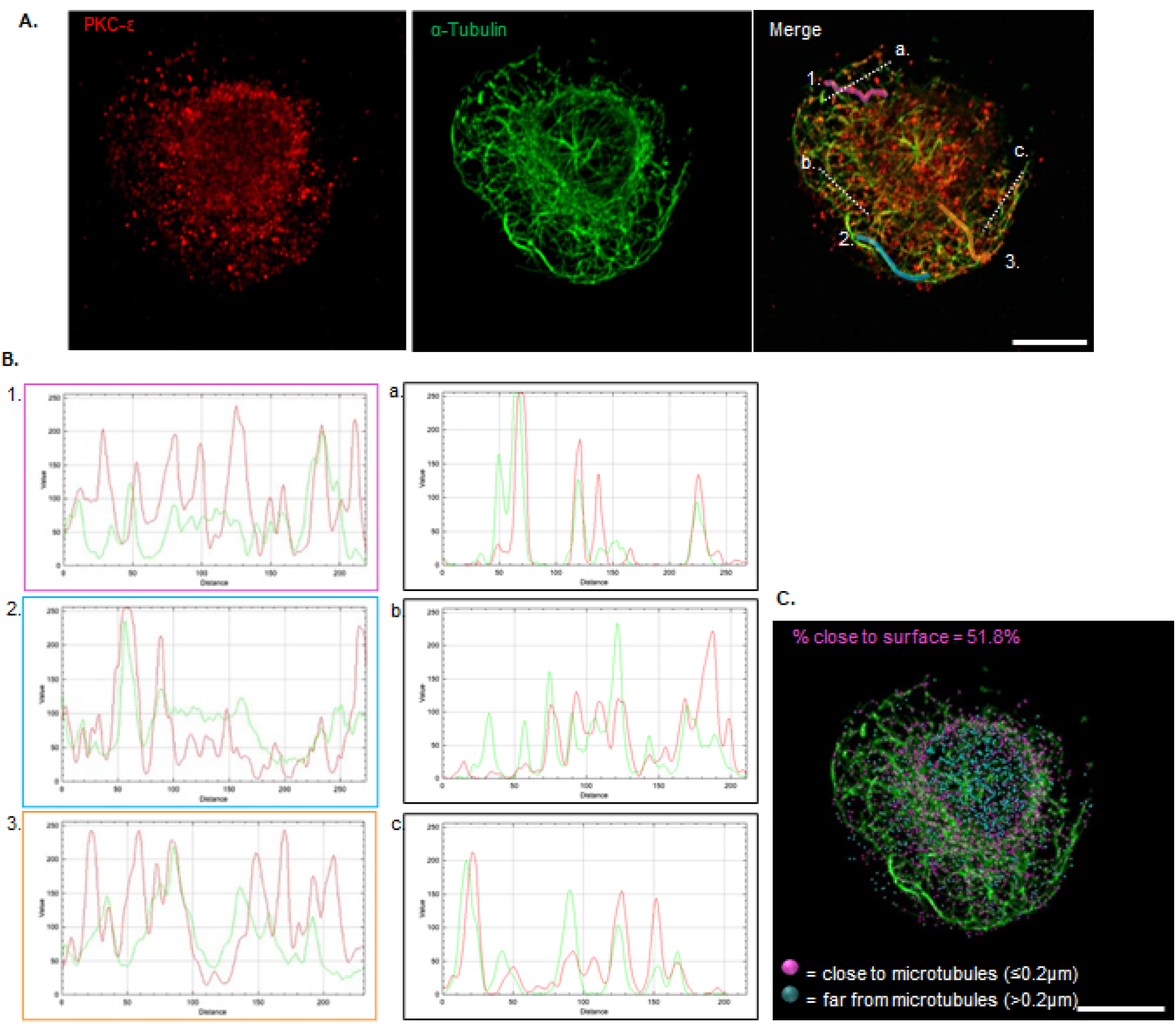
PKC-ε Vesicles Align Along Microtubules. (A) BMDM were subjected to synchronized phagocytosis, spread on IgG surfaces (30 minutes), fixed and stained for endogenous PKC-ε and microtubules (α-tubulin). Z-stacks were collected with Airyscan high resolution microscopy and 3D projections generated with Imaris software. (B) Line plot analysis of PKC-ε and α-tubulin fluorescent intensity alignment along microtubules (1-3) and across microtubules (a-b). Colored borders correspond to numbered lines in merged image (A) and dotted lines correspond to cross-section line plots. (C) 3D b).rendering of PKC-ε puncta from (A). Surfaces were generated from the α-tubulin fluorescence; spots were generated from the PKC-ε fluorescence. Imaris analysis calculated the alignment of PKC-ε spots along the microtubule surface. Aligned PKC-ε puncta (pink spots) were defined as those within 0.2μm of a microtubule, more distant spots were colored blue. 52% of the PKC-ε spots were within 0.2 μm of a microtubule, more distant spots were coloured blue.52 % of the PKC-ε spots were within 0.2 μm of a microtubule (18 cells from 3 independent experiments).

While spreading on IgG-coated surfaces allows visualization of events occurring at the “phagosome”, IgG surfaces do not replicate the focal activation and cross-linking of FcγR that occur during target ingestion. If PKC-ε traffics from the Golgi on microtubules, we would predict that PKC-ε^+^ vesicles preferentially localize on microtubules directed towards the phagosome. To test this, BMDM were incubated with IgG-opsonized targets, fixed and stained for endogenous PKC-ε and α-tubulin (Figure 5A and Supplemental 4A). We quantified the alignment ratio of PKC-ε vesicles along microtubules directed towards the phagosome versus those in non-involved regions of the cell. Briefly, PKC-ε vesicles within 0.2μm of microtubules were defined as “close to/aligned along” microtubules while those further than 0.2μm were designated “far/non-aligned”. Significantly more PKC-ε vesicles were aligned along microtubules directed towards the phagosome compared to microtubules in regions of the cell lacking targets (Figure 5B,C and Supplemental 4B). These data reveal that PKC-ε^+^ vesicles preferentially align along phagosomally-directed microtubules and support a model in which PKC-ε traffics from the Golgi to sites of phagocytosis on vesicles that travel on microtubules.

**Figure 5:**
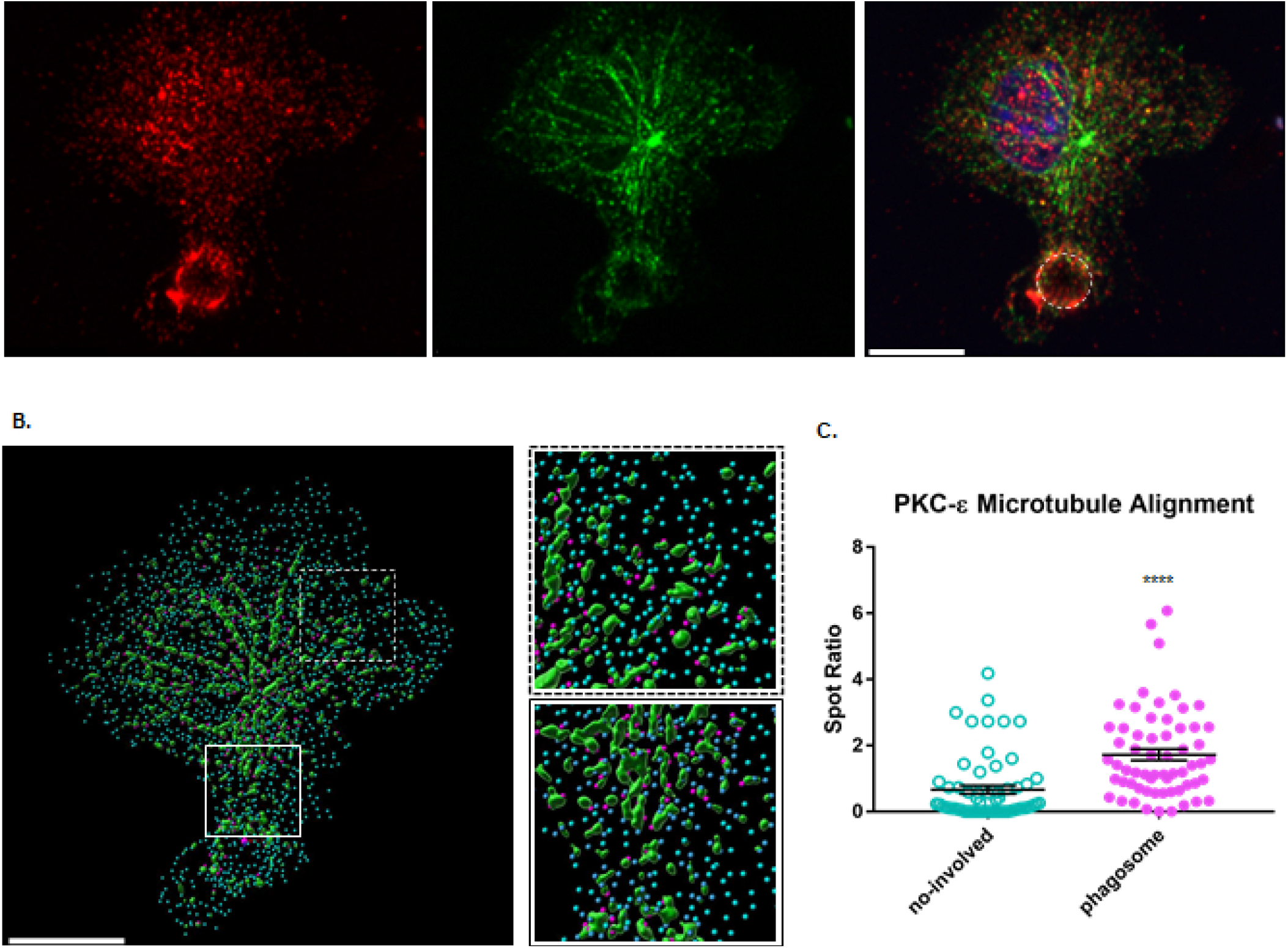
PKC-ε Preferentially Aligns Along Phagosomally Directed Microtubules. BMDM were incubated with 5 μm IgG-coated targets, fixed, and stained for PKC-ε (Red), microtubules (a-tubulin, green), and the nucleus (DAPI). Z-stacks were collected with Airyscan high resolution microscopy. (A) 3D projection of a BMDM during phagocytosis. PKCε, α-tubulin, and the merge image are shown. (B) 3D rendering of z-stacks into PKC-ε spots and microtubule surfaces by Imaris allowed quantitation of aligned PKC-ε spots as detailed in Fig 5. (C) Quantitation of the ratio of aligned (pink) to non-aligned (blue) PKC-ε puncta on microtubules directed towards phagosomes (solid box) and non-involved regions (dotted box) revealed that PKC-ε preferentially aligns on MT directed towards phagosomes. Representative of 57 phagosomal events from 3 independent experiments, each data point represents an event. Statistical significance was determined by paired Student’s t-test.****p<0.0001, Scale bar = 10μm

### PKC-ε Vesicles Contain TNF-α

Colocalization of PKC-ε with VAMP3 (Figure 1) confirms that PKC-ε is on vesicles. If PKC-ε traffics on Golgi-derived vesicles, those vesicles should contain Golgi-associated protein(s). The Stow lab reported that tumor necrosis factor-alpha (TNF-α) traffics from the TGN to the phagocytic cup on vesicles (Manderson et al., 2007; Murray et al., 2005). That TNF-α is synthesized as a transmembrane protein makes it a bona fide marker of Golgi-derived vesicles. Notably, the Golgi-to-phagosome trafficking pattern of TNF-α during phagocytosis parallels that of PKC-ε. Thus, we tested the hypothesis that PKC-ε and TNF-α are transported on the same vesicles. First, RAW cells co-expressing PKC-ε-GFP and TNF-α-mCherry were subjected to synchronized phagocytosis, fixed and imaged (Figure 6A). A Pearson’s correlation coefficient for PKC-ε-GFP and TNF-α-mCherry was calculated at the phagosome, the Golgi, and a non-involved region of the membrane. PKC-ε/TNF-α colocalization at the phagosome and Golgi were significantly higher compared to non-involved regions of the membrane (Figure 6A, graph). Further, BMDM undergoing synchronized phagocytosis were stained for *endogenous* PKC-ε and TNF-α (Figure 6B). High resolution Airyscan again revealed significantly higher colocalization between PKC-ε and TNF-α puncta at the phagosome compared to an equivalent ROI in a region of the call lacking targets (Figure 6B,C). These results, in a cell line and primary macrophages, using exogenous fluorescent TNF-α and PKC-ε or their endogenously stained counterparts, produced the same results: significantly more colocalization of the two markers at the phagosome compared with other parts of the cells, consistent with their Golgi-to-phagosome trafficking on the same vesicles.

**Figure 6:**
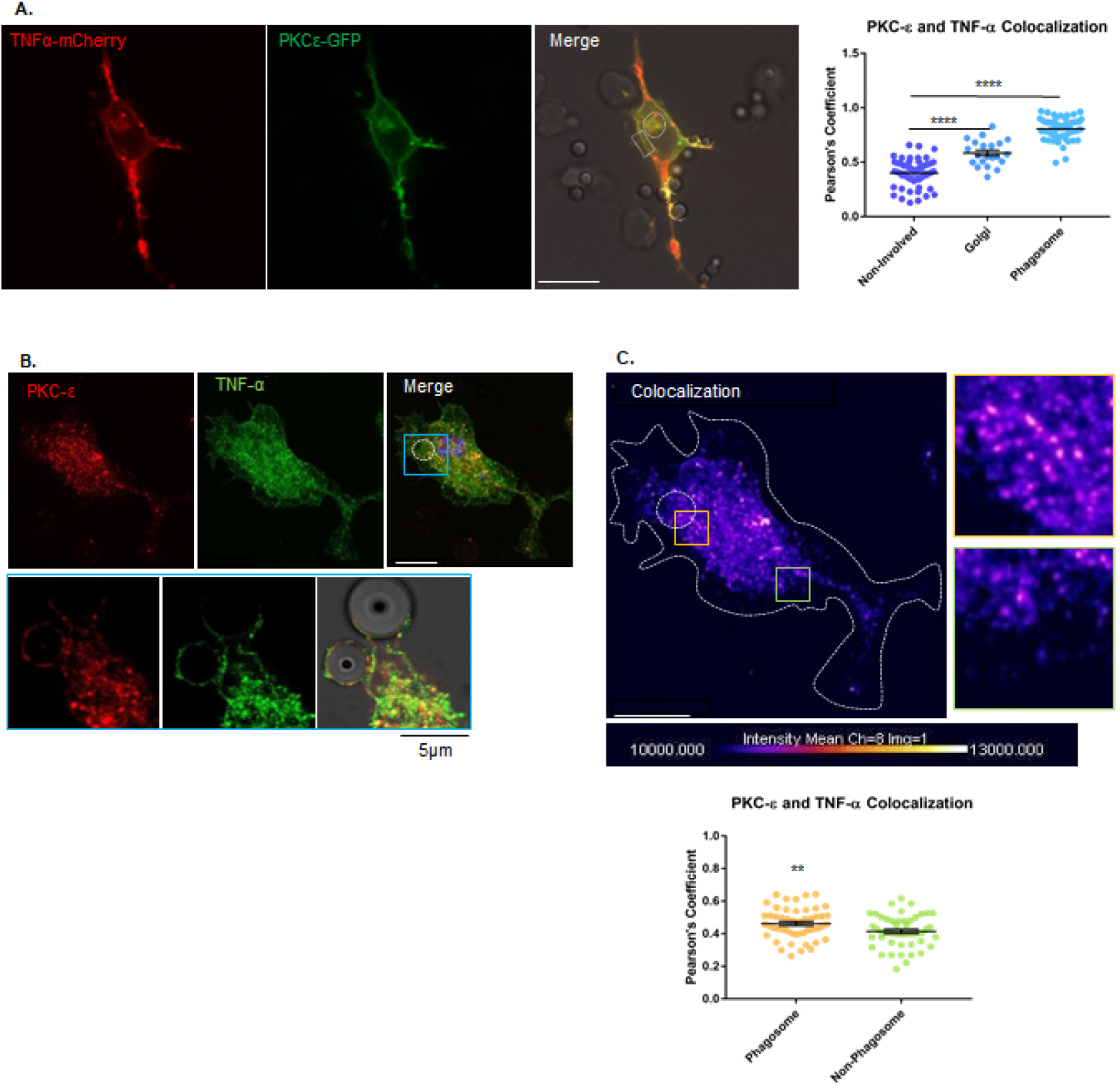
PKC-ε Colocalizes with TNF-a in vesicles and at phagosomes. (A) RAW cells, co-expressing with TNF-α-mCherry and PKC-ε -GFP, were subjected to synchronized phagocytosis, fixed, and z-stacks collected. Images are a single slice from a z-stack. (Graph) Pearson’s correlation coefficient for colocalization between TNF-α and PKC-ε at the Golgi (circle), the phagosome (dotted line), and a region of the plasma membrane not involved in phagocytosis (boxed region). Data are presented as mean ± SEM with each symbol representing one cell. Aggregate data from 4 independent trials (21-40 cells). Statistical significance was determined by one-way ANOVA with Bonferroni post-test. **** p<0.0001. Scale bar = 10μm. (B, Top) BMDM internalizing 5μm IgG-opsonized particles. Cells were fixed and stained for endogenous TNF-α and PKC-ε. Z-stacks were collected with Airyscan high resolution microscopy, images are the 3D projection of z-stack. Scale bar = 10 pm. Boxed region is enlarged below. (Bottom) Single-slice of boxed region showing colocalization of TNF-α and PKC-ε in puncta at the phagosome. Scale bar = 5μm. (C) Colocalization is higher in phagosome than non-phagocytic region of the cell. ROI for phagosome (yellow box) and non-involved (green box) cell regions indicated. LUT represents colocalization in 3D rendered cells, with colocalization indicated on a purple (low) to white (high) color scale. Pearson’s correlation coefficients at these regions were calculated using Imaris software. Quantitation for 51 total cells from 3 independent experiments reveals significantly more colocalization in puncta associated with the phagosome vs non-phagosomal regions. Data are presented as mean ± SEM with each symbol representing one cell. Statistical significance was determined by paired Student’s t-test. ** p<0.01. Scale bar = 10μm.

The high degree of PKC-ε/TNF-α colocalization at the phagosome supports our hypothesis but raises the question: Does PKC-ε associate indiscriminately with intracellular vesicles? To test this, we co-stained BMDM for PKC-ε and either TNF-α, TGN38 (a protein that shuttles between the Golgi and plasma membrane), or EEA1, a marker of early endosomes. Indeed, a calculation of Pearson’s correlation coefficients indicate that PKC-ε colocalizes extensively with TNF-α (0.74, n= 5, 38 cells), modestly with TGN38 (0.48, n= 5, 36 cells) and poorly with the early endosome marker, EEA1 (0.29, n= 5, 30 cells). The high correlation of PKC-ε with TNF-α indicates selective association on Golgi-derived vesicles. Its modest association with TGN38 is likely due to the fact that TGN38 cycles from the TGN to the plasma membrane. PKC-ε likely associates with the exocytic, but not endocytic, phase of that cycle. The low association with EEA1 would be consistent with this interpretation.

### Delivery of Vesicles to the Phagosome Requires PKC-ε

We know that dissociation of PKC-ε from the Golgi 1) inhibits its concentration at the phagosome, 2) blocks membrane addition in response to FcγR ligation, and 3) significantly reduces phagocytosis (Hanes et al., 2017). Using TNF-α as a surrogate marker for Golgi-derived vesicles, we asked if PKC-ε is necessary for delivery of TNF-α to the phagosome. We reasoned that, if PKC-ε and TNF-α are transported on the same vesicles, and Golgi-tethered PKC-ε is necessary for their delivery, then dissociation of PKC-ε from the TGN (by expression of GFP-hSac1-K2A or treatment with PIK93) should inhibit TNF-α concentration at the phagosome. Indeed, expression of GFP-hSac1-K2A in RAW cells significantly reduced TNF-α concentration at the phagocytic cup (Figure 7A). Similarly, WT BMDM treated with PIK93 delivered significantly fewer TNF-α puncta to the phagosome compared with controls (Figure 7B). However, TNF-α delivery is restored upon PIK93 washout or when PKC-ε is expressed in εKO cells (Figure 7B). Notably, TNF-α colocalizes with PKC-ε puncta in both WT and εKO re-expressing BMDM (Figure 7C). Together, these data provide a third line of evidence that PKC-ε is required for TNF-α vesicle delivery.

**Figure 7:**
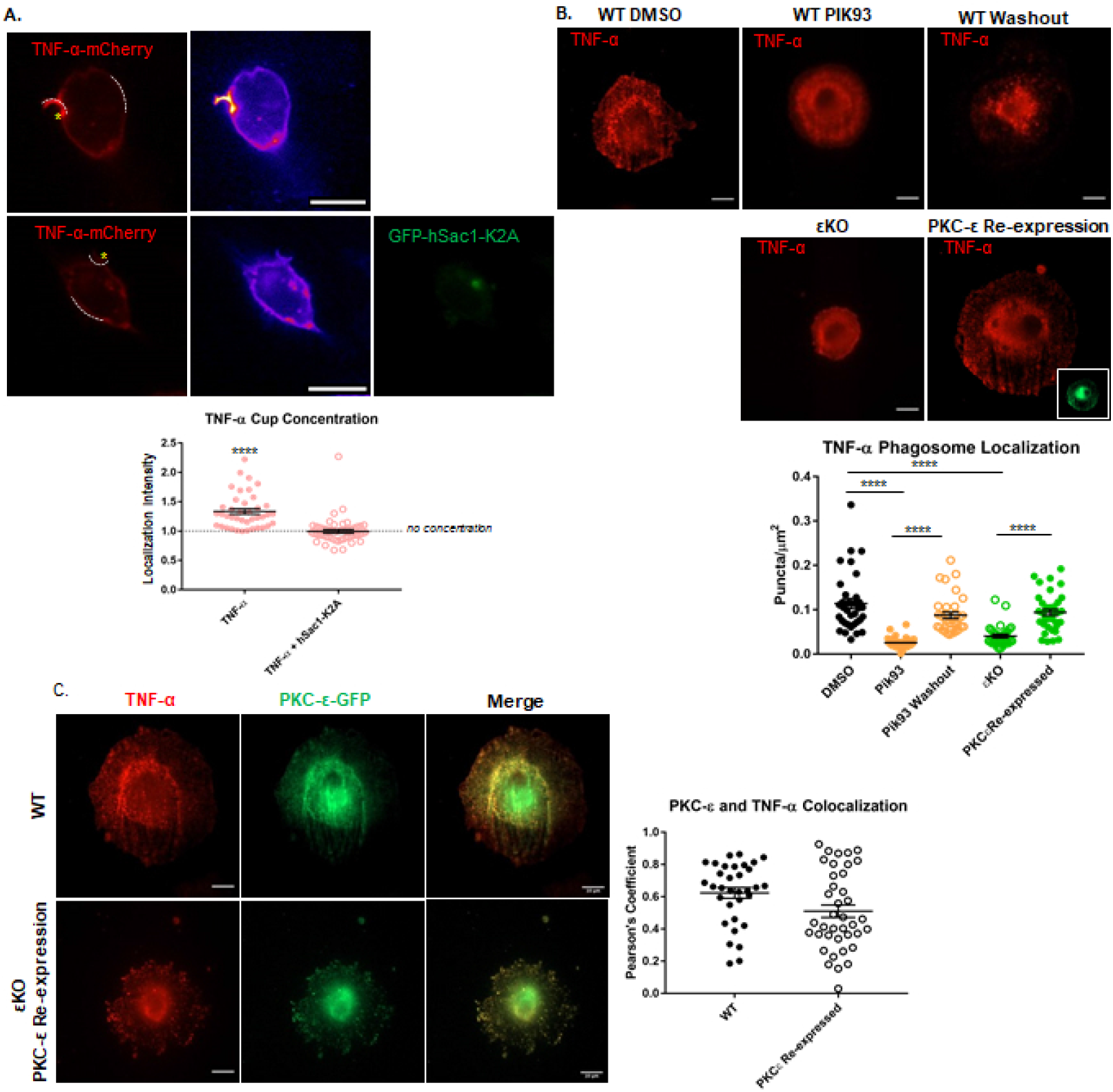
Golgi-tethered PKC-ε is necessary for TNF-α concentration at phagosomes. (A) RAW cells were transfected with TNF-α-mCherry ± GFP-hSac1-K2A and subjected to synchronized phagocytosis. Z-stacks were collected by spinning-disc confocal microscopy. The localization intensity of TNF-α is reported as the intensity at the cup normalized to an equivalent membrane ROI in a non-involved region of the cell (indicted with dotted lines). TNF-α concentration at phagocytic cups was significantly reduced in cells co-expressing GFP-hSac1-K2A (bottom panel). (Middle column) Pseudocolored LUT was applied for optimal visualization of TNF-α fluorescent intensity. GFP-hSac1-K2A localizes to the Golgi, depleting Golgi-tethered PKC-ε. Data are presented as mean ± SEM from 3 independent experiments, (40-51 phagosomes), each data point represents the normalized density at a phagosome. Statistical significance was determined by unpaired Student’s t-test. **** p<0.0001. Scale bar = 10μm. (B) BMDM from WT and PKC-ε null (ε KO) mice were spread on IgG surfaces, fixed, stained for endogenous TNF-α, and imaged by TIRFM. Some WT cells were treated with PIK93 (100 nM, 30 min) and either fixed immediately or the drug was washed out (5 min) prior to fixation. PKC-ε-GFP was expressed in ε KO BMDM to determine how its re-expression affects TNF-α delivery. Left, representative images of 34-37 cells/condition form 3 independent experiments. Right, quantitation of TNF-α puncta, normalized to cell area; each data point represents a single cell. Data are presented as mean SEM. Statistical significance was determined by one-way ANOVA with Bonferroni post-test. **** p<0.0001. DMSO VS PKC-ε re-expression is n.s. PIK93 vs ε KO is n.s. Scale bar = 10μm. (C) PKC-ε-GFP expressed in WT and ε KO BMDM. Cells were spread on IgG surfaces and stained for endogenous TNF-O.. Representative images are shown (left). (Right) significant colocalization between PKC-ε and TNF-α was determined by Pearson’s correlation coefficient (graph). Data are presented as mean ±SEM. n = 3 (27-39 cells, per condition) with each data point representing a cell. Statistical significance was determined by unpaired Student’s t-test. Colocalization in WT and PKC-ε re-expressing εko cells was not statistically different. Scale bar = 10μm.

### The Regulatory Domain of PKC-ε is Sufficient for Vesicle Delivery

Our earlier work revealed that the isolated regulatory domain of PKC-ε (εRD, aa 1-900) concentrates at the phagocytic cup (Cheeseman et al., 2006; Wood et al., 2013). Given that knowledge, and the current results suggesting that PKC-ε orchestrates TNF-α delivery, we asked if εRD *per se* supports vesicle transport. Using TIRFM and frustrated phagocytosis, we quantified TNF-α puncta at the membrane in WT and εKO BMDM expressing εRD-GFP. The number of εRD puncta in WT and εKO were not significantly different from each other or from WT BMDM expressing full-length PKC-ε-GFP (Figure 8A). Interestingly, while TNF-α vesicles are not delivered in εKO BMDM (Figure 7B), they *are* upon expression of εRD in these cells (Figure 8B,C) and TNF-α colocalizes with εRD (Figure 8B,D). These results uncover a novel and intriguing pathway in which the regulatory domain of PKC-ε, independent of its catalytic domain/activity, is necessary for the delivery of Golgi-derived vesicles to the phagosome. As we have previously shown that PKC-ε’s catalytic activity is necessary for membrane fusion (Hanes et al, 2017), these findings are consistent with a model in which PKC-ε serves two functions during phagocytosis: εRD drives vesicle delivery while catalytic activity phosphorylates substrates required for vesicle fusion. These findings provide a foundation for further exploration into the role of εRD at the Golgi and identification of PKC-ε substrates at the phagosome.

**Figure 8:**
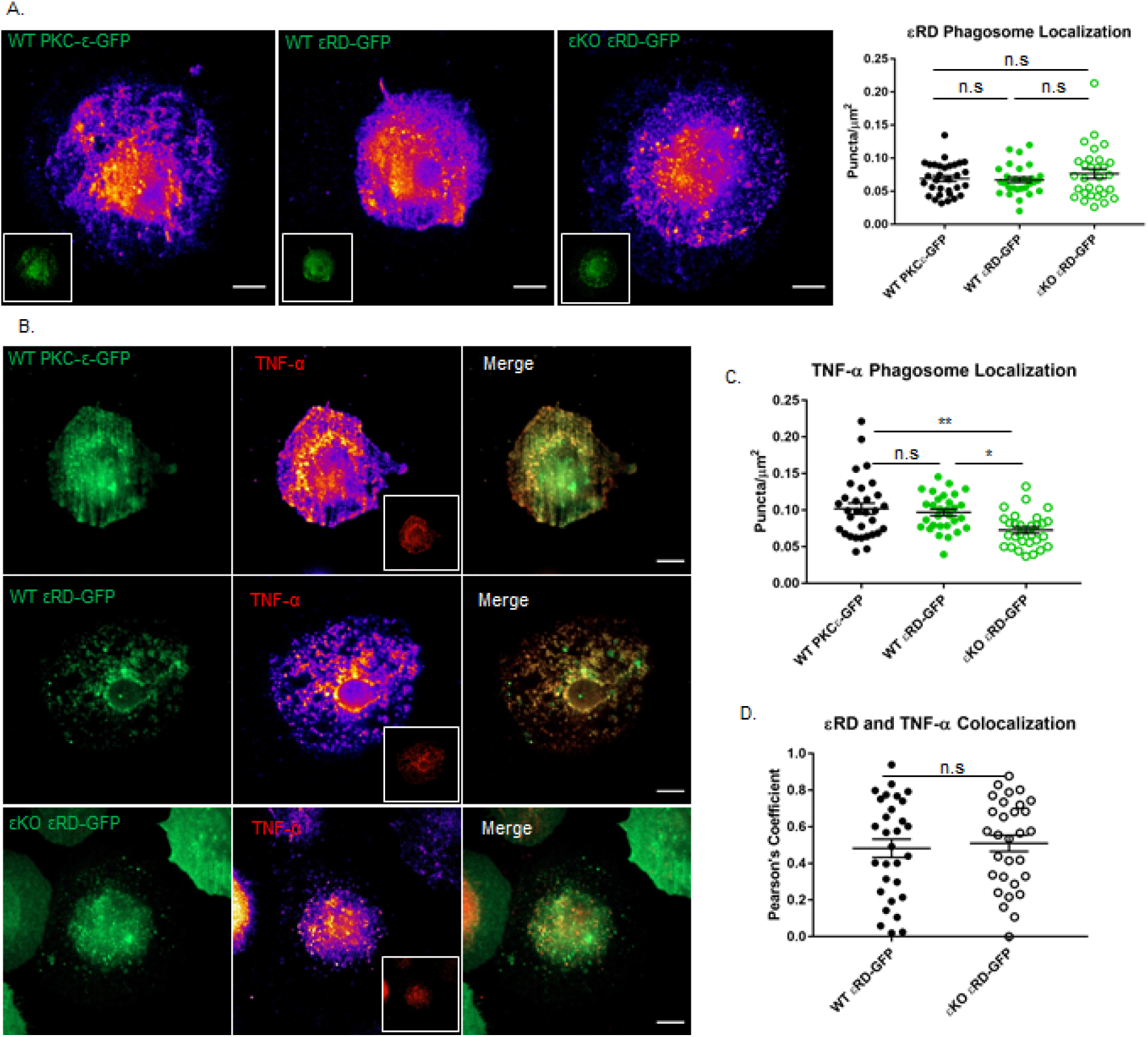
The Regulatory Domain of PKC-ε is Sufficient for TNF-α Vesicle Delivery. (A,B) WT and εKO BMDM, expressing either PKC-ε-GFP or the regulatory domain of PKC-ε (εRD-GFP) were spread on IgG surfaces, fixed, stained for endogenous TNF-α and imaged by TIRFM. (A) PKC-ε-GFP and εRD-GFP expression in WT and εKO BMDM. Representative images are shown with puncta visualized with a Fire LUT with corresponding GFP in inserts. (A, graph) The number of puncta/cell area was calculated as in Fig 3. Data are presented as mean ± SEM with each datapoint representing one cell. n = 3 independent experiments with a total of 30-31 cells per condition. Statistical significance was determined by One-way ANOVA with Bonferroni post-test. There was no significant difference in the number of PKC-ε-GFP or εRD-GFP puncta. (B) TIRF images of endogenous TNF-α staining in PKC-ε-GFP and εRD-GFP expressing BMDM. Fire LUT was applied for optimal visualization of TNF-α fluorescence and puncta, with corresponding TNF-α fluorescence staining in inserts. (C) The number of TNF-αpuncta/cell area was calculated the same as in above and Figure 3. Data are presented as mean ±SEM. n = 3 (30 cells per condition); each data point represents a cell. Statistical significance was determined by Oneway ANOVA with Bonferroni post-test. **p<0.01, *p<0.05. There was no significant difference in TNF-α puncta in WT BMDM expressing PKC-ε-GFP and εRD-GFP. (D) Pearson’s correlation coefficient between endogenous TNF-α and εRD-GFP was quantitated. Data are presented as mean ±SEM. n = 3 (30 cells per condition), each data point represents a cell. Statistical significance was determined by paired Student’s t-test. There was no significant different in colocalization between TNF-α and εRD-GFP. Scale bars = 10 μm.

In summary, these studies reveal a novel paradigm for focal exocytosis in non-polarized cells. Specifically, PKC-ε translocates on vesicles that travel on microtubules. The vesicles carry TNF-α from the Golgi to the phagosome and PKC-ε is required for their delivery. The vesicles fuse, expanding the membrane for pseudopod extension and rapid phagocytosis. The most intriguing finding from this work is that PKC-ε’s regulatory domain is required for the transit of vesicles, defining a PKC function independent of its catalytic activity. While many questions remain to be answered, these findings open the door to investigations of PKC-ε orchestration of focal exocytosis in non-polarized cells.

## Discussion

Efficient IgG-mediated phagocytosis requires membrane addition for pseudopod extension (Braun and Niedergang, 2006; Cannon and Swanson, 1992; Gerlach et al., 2020; Holevinsky and Nelson, 1998) but the source of that membrane and the mechanism for its focal delivery is unclear. Work by the Grinstein lab demonstrated the targeted delivery of VAMP3^+^ vesicles to forming phagosomes (Bajno et al., 2000). Their findings unequivocally demonstrated focal exocytosis of vesicles at forming phagosomes. Our studies identified PKC-ε as a critical player for pseudopod extension and whole cell patch clamping established that macrophages expand their membrane by ∼1/3 as they spread on IgG surfaces (Hanes et al., 2017; Wood et al., 2013). PKC-ε null macrophages were unable to add membrane is response to FcγR ligation, a deficit that was reversed upon expression of PKC-ε in PKC-ε null BMDM (Hanes et al., 2017).

As the majority of macrophage PKC-ε is cytosolic (Larsen et al., 2000), and PKCs translocate from the cytosol to their sites of activity, we predicted that cytosolic PKC-ε concentrated at phagosomes to facilitate membrane addition. Several findings led us to rethink this assumption. First, using capacitance to quantitate membrane expansion, we found no addition of membrane in cells treated with the microtubule disrupting drug nocodazole, or with PIK93 which selectively dissociates PKC-ε from the Golgi (Hanes et al., 2017). If there was significant cytosolic translocation, we would have expected at least some membrane addition in the presence of these inhibitors. Secondly, our previous studies revealed a Golgi-associated pool of PKC-ε. We discovered that the pseudosubstrate region of PKC-ε interacts with PI4P, tethering PKC-ε to the TGN. Blocking this interaction through selective removal of PI4P with hSac1-K2A, or inhibition of PI4 Kinase with PIK93, dissociates PKC-ε from the TGN, prevents PKC-ε concentration at phagosomes, and significantly reduces phagocytosis (Hanes et al., 2017). These findings support a model in which the Golgi-tethered, rather than cytosolic, PKC-ε is critical for phagocytosis. Notably, hSac1-K2A and PIK93 provided us with tools to selectively deplete the Golgi-tethered pool of PKC-ε, allowing us to study its role in phagocytosis. Finally, during live imaging of phagocytosis in PKC-ε-GFP expressing BMDM, we observed PKC-ε puncta exiting the Golgi (Video 1), providing real time evidence that PKC-ε translocates from the Golgi on a vesicle, an unexpected discovery.

This raised the question: How is Golgi-tethered PKC-ε transported to the phagosome? The most likely mechanism would be vesicular trafficking on microtubules. Indeed, others have reported that delivery of lipids and proteins from the Golgi to the plasma membrane on vesicles (Stalder and Gershlick, 2020). We had reported that nocodazole prevented membrane addition in response to FcγR ligation (Hanes et al, 2017), implicating microtubules and, by extension, vesicular trafficking in phagosome formation. TIRFM allowed us to visualize the “phagosome”, revealing that PKC-ε approaches the membrane on a vesicle, a vesicle that then fuses (Figures 1-3). To our knowledge, this is the first evidence that *any* PKC translocates on a vesicle to the plasma membrane and represents the first novel finding of this work.

If Golgi-tethered PKC-ε traffics on vesicles, but its dissociation from the Golgi blocks phagocytosis and PKC-ε concentration (Hanes et al., 2017), how could we study the role of PKC-ε at the Golgi? Work from the Stow lab documenting that TNF-α (a transmembrane protein that travels through the Golgi) concentrates at phagosomes (Manderson et al., 2007; Murray et al., 2005) provided us with a candidate marker for phagosomally directed, Golgi-derived vesicles. Once we established that PKC-ε and TNF-α colocalized at the Golgi and at the phagosome (Figure 6), we could use TNF-α to ask if PKC-ε was simply a passenger on vesicles or played a more active role. Quantifying TNF-α appearance at the membrane in PKC-ε null BMDM, and upon re-expression of PKC-ε in null cells, revealed that PKC-ε is required for TNF-α^+^ vesicle trafficking to the phagosome, representing a second novel finding of this work: that PKC-ε is required for Golgi-derived vesicle delivery to the sites of FcγR ligation membrane (Figure 7).

Given that PKC-ε transits from the Golgi to phagosomes on vesicles along microtubules and drives vesicle delivery, we are left with the question: *How* does PKC-ε facilitate Golgi-to-phagosome vesicular trafficking? PKC-ε’s catalytic activity is required for membrane fusion, but not for concentration of PKC-ε at the phagosome (Hanes et al., 2017; Wood et al., 2013). The regulatory domain of PKC-ε concentrates at the phagosome, but is it sufficient for delivery of vesicles? εRD-GFP expressed in εKO macrophages appeared as puncta by TIRFM, indistinguishable from the pattern produced by full-length PKC-ε-GFP (Figure 8A). That expression of εRD-GFP in εKO cells restored delivery of TNF-α puncta demonstrating that εRD is sufficient for vesicular trafficking from the Golgi (Figure 8B), the third novel finding from this work.

Finally, what, if any, is the role of VAMP3 during this process? VAMP3 is a marker of recycling endosomes (McMahon et al., 1993). While it colocalizes with TNF-α and PKC-ε at the phagosome (Figure 2B), it is not required for FcγR-mediated phagocytosis (Allen et al., 2002). One model that is consistent with all the data would be that PKC-ε (actually εRD) orchestrates vesicle formation at the Golgi (D’Amico and Lennartz, 2018; Miralles, 2018). The vesicles then “mature” as they move to phagosome, fusing with VAMP3^+^ endosomes before or when docking at the phagosome. Upon docking, the catalytic activity of PKC-ε phosphorylates as yet unknown substrate(s) for vesicle fusion, pseudopod extension and phagocytosis.

### Summary

The data presented include three novel findings: 1) PKC-ε is delivered to the phagosome on vesicles, 2) these vesicles originate in the Golgi, carry TNF-α, and their delivery requires PKC-ε, and 3) the regulatory domain of PKC-ε is sufficient for vesicle delivery. These findings challenge the current paradigm of PKC activation by translocation from the cytosol and raise numerous questions, the answers to which will provide insight into novel PKC functions. Unanswered questions include: What is the role of PKC-ε at the Golgi? How does the regulatory domain, independent of the catalytic domain (and activity) mediate vesicular trafficking? How are vesicles targeted to the phagosome? And what is/are the PKC-ε substrates at the phagosome that are required for vesicle fusion?

Perhaps the most exciting avenue of investigation will be the relay between FcγR ligation and the Golgi. Pilot studies indicate that Syk signaling is required for PKC-ε concentration at the phagosome. If so, then there must be some FcγR-to-Golgi-to-phagosome relay system that rapidly recruits Golgi vesicles specifically to phagosomes for focal exocytosis. Elucidation of this signaling network may provide insight into diseases such as cancer, where elevated PKC-ε promotes focal exocytosis for tumor cell metastasis (Gorin and Pan, 2009; Pan et al., 2005; Tachado et al., 2002).

## Materials and Methods

### Buffers and Reagents

ACK Lysis Buffer: 0.5 M NH_4_Cl, 1 mM KHCO_3_, and 0.1 mM EDTA (pH 7.4). HBSS^++^: HBSS containing 4 mM sodium bicarbonate, 10 mM HEPES, 1.5 mM each CaCl_2_, and MgCl_2_ (pH 7.4). Paraflormaldehyde (PFA) Polysciences, Inc (18814-10). DMSO (Sigma, D2438). PIK93 was from Echelon Biosciences (Salt Lake City, Utah) used at 100nM. Nocodazole was from Cayman Chemicals (Ann arbor, Michigan) used at 10μM. Blocking buffer: PBS containing 0.5% fish gelatin, 2% BSA, and thimerosol.

### Mice and Cells

PKC-ε^+/-^heterozygotes on the C57/B16 background were purchased from the Jackson Laboratory (stock# 004189; Bar Harbon, ME) and bred in the Albany Medical Center Animal Resources Facility. Heterozygotes were crossed to produce the wild type and PKC-ε^-/-^mice. All animal procedures were approved by the Albany Medical Center Institutional Animal Care and Use Committee. Cells obtained from one mouse represent one independent experiment.

#### Bone marrow-derived macrophages (BMDM)

PKC-ε^+/+^ and PKC-ε^-/-^mice were euthanized and their femurs and pelvis were removed. Bone marrow was collected, and red blood cells were lysed with ACK lysis buffer. Bone marrow stem cells were differentiated by incubation in bone marrow media (BMMM) containing phenol-red free (high glucose) DMEM, 10% PBS, 20% L-cell conditioned media, sodium bicarbonate, and gentamicin (RPI, G38000). BMDM were used 7-14 days after harvesting.

#### RAW 264.7 mouse macrophages

The RAW subclone, LacR/FMLPR.2 (Raw cells) were used. Cells were maintained in media containing DMEM, 10% FBS, and sodium bicarbonate.

### IgG-Coating Targets and Surfaces

Acid-washed 2 or 5μm boroscilicate glass beads (Duke Standards, Thermo Scientific, 9005) or 15mm glass coverslips/matteks were sequentially coated with poly-L-lysine, dimethylpimelimidate, and 1% IgG-free BSA. Free active groups were blocked with 0.5M Tris (pH 8.0) (1 hour, room temperature) before opsonizing with rabbit anti-BSA IgG (Sigma, B1520). IgG-free BSA treatment and blocking prior to opsonization was omitted for beads and surfaces opsonized with human IgG (Sigma, I4506). Free active groups were blocked overnight with 0.5M Tris (pH 8.0) (4°C, overnight). Before use, beads and surfaces were washed with PBS and diluted in HBSS^++^.

### Phagocytosis

#### Phagocytosis of IgG-opsonized targets

Time courses with 5μm human IgG-opsonized beads were performed as previously described (Wood et al., 2013). 5μm opsonized beads were added (∼3 beads per cell) and then incubated at 37°C. Cells were fixed (4%PFA/PBS) and prepared for immunofluorescent staining.

#### Frustrated Phagocytosis

Synchronized time courses on human IgG-coated coverslips were done. Cells were cooled on ice for 30 minutes and transferred to a 37°C heating block. Cells were fixed (4%PFA/PBS) after 30 minutes and prepared for immunofluorescent staining. For live imaging, ice steps were eliminated. Cells were imaged via TIRF microscopy as they sat and spread onto rabbit or human IgG-coated surfaces.

#### Synchronized Phagocytosis

Cells were pre-incubated on ice for 30 minutes and then transferred to 37°C heating block. Cells were then treated as noted.

### Expression of Exogenous Proteins

#### Nucleofection

Plasmids Akt-PH-GFP, VAMP-3-GFP, and PKC-ε-GFP were delivered BMDM following manufacturers’ instructions. Mouse macrophage kit (VPA-1009). 3 × 10^6^ BMDM were suspended briefly in 82μL macrophage solution and 18μL supplement. DNA (5μg) was added and cells were transferred to a sterile electroporation cuvette. Nucleofector was set to Y-001 program and after cells were transferred to a 6-well dish with 2mL of BMM.

#### Transfection

RAW 264.7 cells were cultured as described. Cells (5 × 10^4^) were seeded onto 12-mm matteks or coverslips and transfected with Lipofectamine2000 at a 3:1 (Lipofectamine2000:DNA). A detailed protocol has been published (Wood et al., 2013). Akt-PH-GFP, GFP-hSac-1-K2A and TNFα-mCherry plasmids were generous gifts from T. Balla (Varnai and Balla, 1998) P. Mayinger (Mayinger, 2009) and J. Stow (Murray et al., 2005), respectively. Constructions of PKC-ε-GFP plasmid has been previously described (Shirai et al., 1998)

#### Viral Transduction

Retroviral construction of PKC-ε-GFP and BMDM transduction have been previously published (Hanes et al., 2017; Wood et al., 2013).

### Immunofluorescent Staining

#### Antibodies

Mouse anti-PKC-ε (Santa Cruz, sc-1681, 1:50, used only in Figure 1) Mouse anti-TNF-α (abcam, ab1793, 1:100). Bunny anti-PKC-ε (Millipore, 06-991, 1:200). Mouse anti-GM130 (BDBiosciences, bd610822, 1:500). Primary antibodies were diluted in blocking buffer. Anti-α-tubulin-FITC (Sigma-Aldrich, F2168, 1:250 diluted in 1%BSA). Anti-Golgin-245 Alexa555 (Bioss, bs13487R-A555, 1:100 diluted in 1% BSA). Goat anti-bunny Alexa488 (Life Technologies, A21069, 1:500). Goat anti-mouse Alexa488 (Life Technologies, A11017, 1:500) Goat anti-mouse Alexa568 (Life Technologies, A11019, 1:500). Secondary antibodies were incubated in blocking buffer containing 10% serum.

Cells were fixed with 4%PFA/PBS and permeabilized with 0.04%Triton/PBS. Blocking was done for 1 hour at room temperature with blocking buffer. Primary antibodies were incubated overnight, 4°C. Secondary antibodies were incubated for 1 hour, room temperature. Cells were stained with DAPI (5 minutes) and post fixed (1%PFA/PBS, 10 minutes). Coverslips were mounted with Prolong Glass Diamond Mount Media (Invitrogen, P36980) and matteks were stored in thimerosal/HBSS^++^ (4°C).

### Imaging

#### Total Internal Reflection Fluorescence Microscopy

Zeiss laser TIRF 3 system using 100x/1.25 oil objective. Live images were collected every 8 seconds over a 10-minute period.

#### High-Resolution Airyscan Microscopy

Zeiss LSM880 confocal microscope system with Airyscan detector running under Zeiss ZEN2.3 software. A 63x/1.4NA oil objective was used to collect z-stacks and live images. Raw images were processed using airyscan processing in ZEN Black 2.3 software.

#### Spinning-Disk Confocal Microscopy

Olympus IX81-DSU with 100x/1.4N.A. oil objective; Hamamatsu electron-multiplying charge-coupled device camera, driven by MetaMorph software (Molecular Devices, Sunnyvale, CA). Live images were collected every 5 seconds over a 10-minute period.

### Analysis

#### Vesicle Fusion

Videos were analyzed using FIJI software. Fusing vesicles were isolated by cropping and the maximum and minimum fluorescent intensities were measured and plotted. Measurements were taken each frame, starting before the vesicle appears to after the fusion event. Intensities for each event were plotted onto separate graphs. Pseudo-colored LUT was applied to images to better visualize fusion events. 3D surface plots and frame montage were also generated in FIJI.

#### Microtubule Alignment

FIJI software was used to generate color line plots of DM1-α and PKC-ε fluorescent intensities. For 3D reconstructions, surfaces were generated based on DM1-α staining and spots for PKC-ε puncta in Imaris x64 version 9.5.0. Xtension “find spots close to surface” was performed with a threshold set to 0.2μm. Percentage of spots within 0.2μm or closer (“close spots”) was quantitated in cells undergoing frustrated phagocytosis. ROIs for cells phagocytosing targets, ROIs of 6μm by 6μm around the phagosome or a non-involved region of the cell were generated. The number of “close” PKC-ε spots to DM1-α surface was divided by the number of PKC-ε puncta further than 0.2μm (“far spots”) to generate a ratio. Ratio of close versus far spots was analyzed in Prism.

#### Phagosome Co-Localization

Imarisx64 version 9.5.0. 3D renderings of phagosome were created based on TNF-α or PKC-ε fluorescence expression. Mask of 3D rendering was applied to fluorescent channels. Pearson’s correlation coefficient was calculated. ROIs for co-localization were made from masked channels. Same steps were applied to a non-involved region of the membrane for comparison.

#### Vesicle Co-localization

Imarisx64 version 9.5.0. For airyscan images, 3D renderings were created in phagosome and non-involved ROIs based on fluorescent stains and used to mask each channel. Pearson’s correlation coefficient was calculated between masked PKC-ε and TNF-α channels. For TIRF images, background was subtracted, and a mask ROI was applied based on PKC-ε staining. Colocalization analysis of phagocytic and non-involved regions was calculated using Pearson’s correlation in Imaris software.

#### TNF-α Concentration at Phagosomes

Analysis of TNF-α at phagosomes was performed using FIJI software (Wood et al., 2013). Mean fluorescence intensity at phagosomes and non-involved regions of similar size. Concentration index was calculated by normalizing phagosome intensities to non-involved regions. Values higher than 1.0 signify concentration.

#### Puncta Quantitation

Find maxima function in FIJI software was utilized to quantitate the number of puncta at the cell surface in TIRF images. Number of puncta was normalized to cell area. Background subtraction was applied to images for optimal visualization of puncta.

#### Statistical Analysis

All data are represented as mean ± SEM. Significance was calculated by paired and unpaired student’s t-test or one-way ANOVA with a Bonferroni post-test. p ≤ 0.05 was considered significant.

## Acknowledgments

We thank Dr. Joseph Mazurkiewicz and the Albany Medical College Imaging Facility for use of the TIRF and high-resolution Airyscan confocal microscopes, Drs. Kate Tubbesing and Ling Wang for assistance with Imaris imaging analysis. Additionally, we would like to thank Drs. Margarida Barroso, Jeremy Logue, and James Drake for constructive feedback and reading the manuscript. Finally, many thanks to Deborah Moran and Rosemary Prestipino for administrative help. Supported by the Johnathan R. Vasilious Foundation.

## Author Contributions

AED conducted and imaged the majority of the experiments and analysis of data and did much of the writing. ACW ran and generated Fig 2B,C and provided movies for Fig 3. CMZ was involved in the planning and protocol development and edited the manuscript. AM conducted experiments in Fig6A, Fig7A, and Supplemental Fig1. MRL supervised all aspects of the study, produced the images for Fig 6A and co-wrote the manuscript.

## List of Abbreviations

PKC-ε: Protein Kinase C-epsilon
εRD: Regulatory domain of PKC-ε
TNF-α: Tumor necrosis factor-alpha
VAMP3: Vesicle associated membrane protein 3
TIRFM: Total internal reflection microscopy
TGN: Trans-Golgi network
FcγR: Fc-gamma receptor
PI4P: Phosphoinositide 4-phosphate
IgG: Immunoglobulin G
Akt-PH: PH domain of Akt

---

**Video 1:**
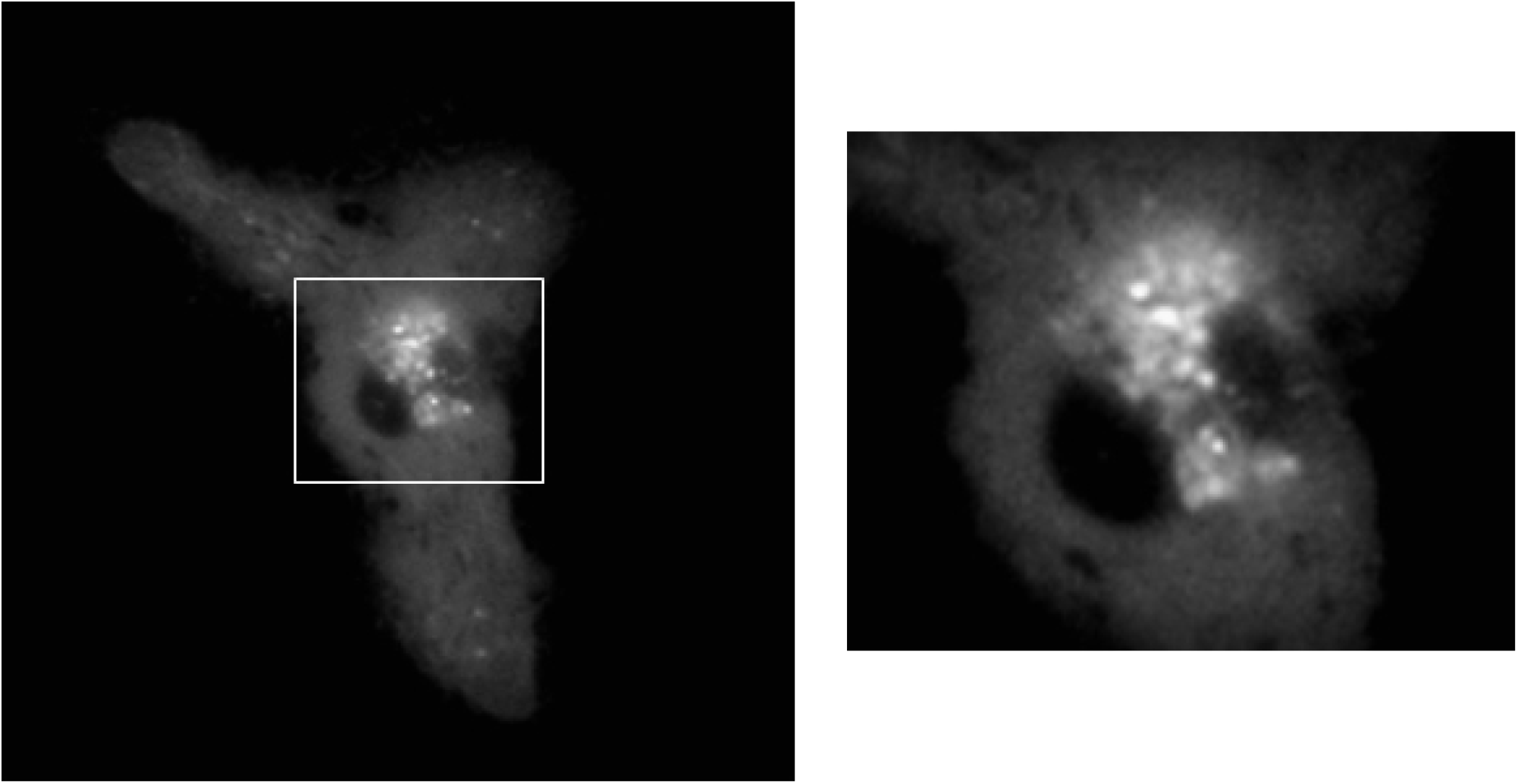
PKC-E Puncta Exiting Golgi. BMDM expressing PKC-ε-GFP phagocytosing IgG-opsonized targets. Images were collected every 3 seconds over a 10-minute period with spinning-disc confocal microscopy

**Video 2:**
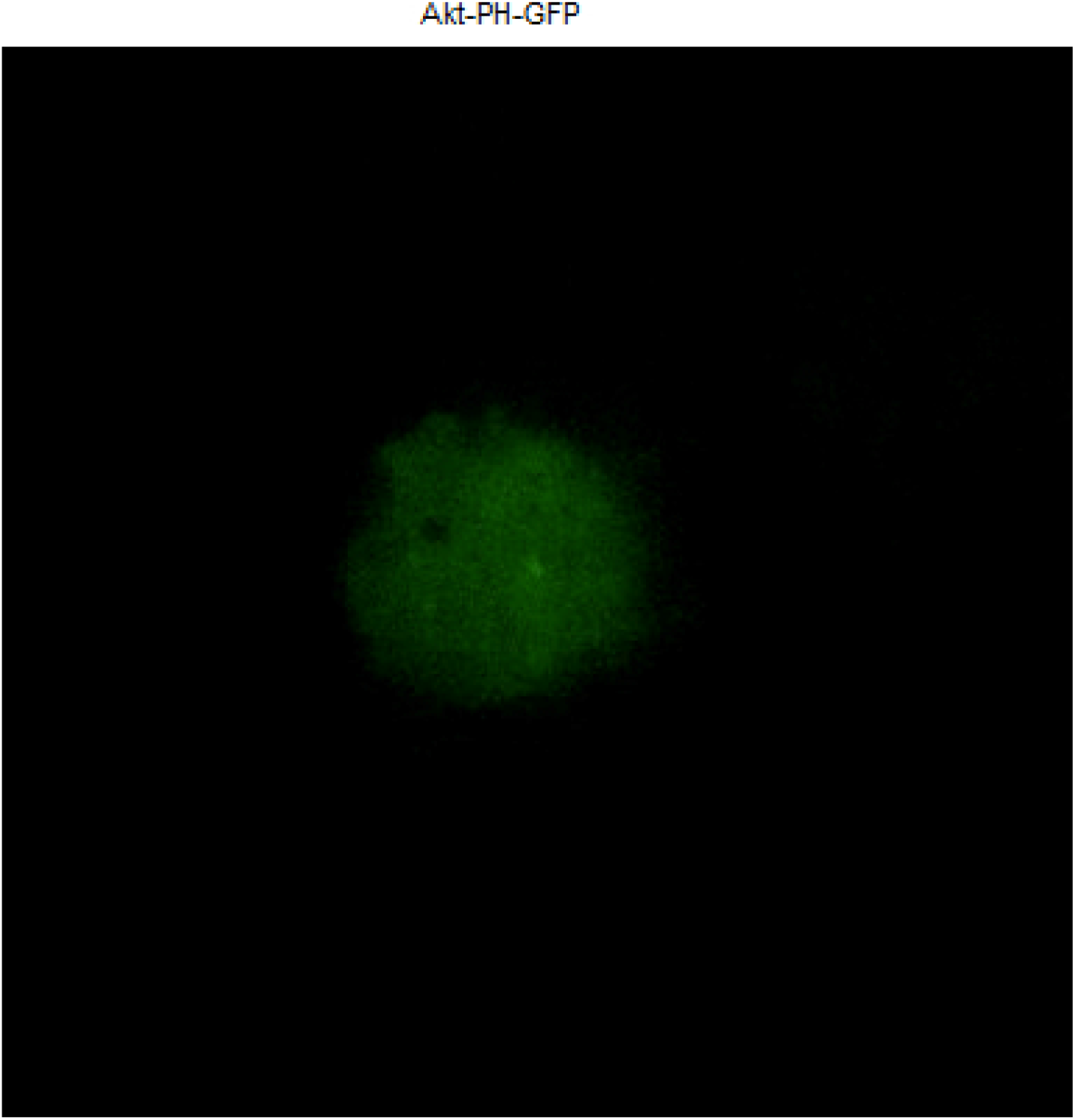
BMDM expressing Akt-PH-GFP were followed with time as the cells spread on IgG surfaces, Images were collected every 8 seconds over a 10-minute period with TIRF microscopy

**Video 3:**
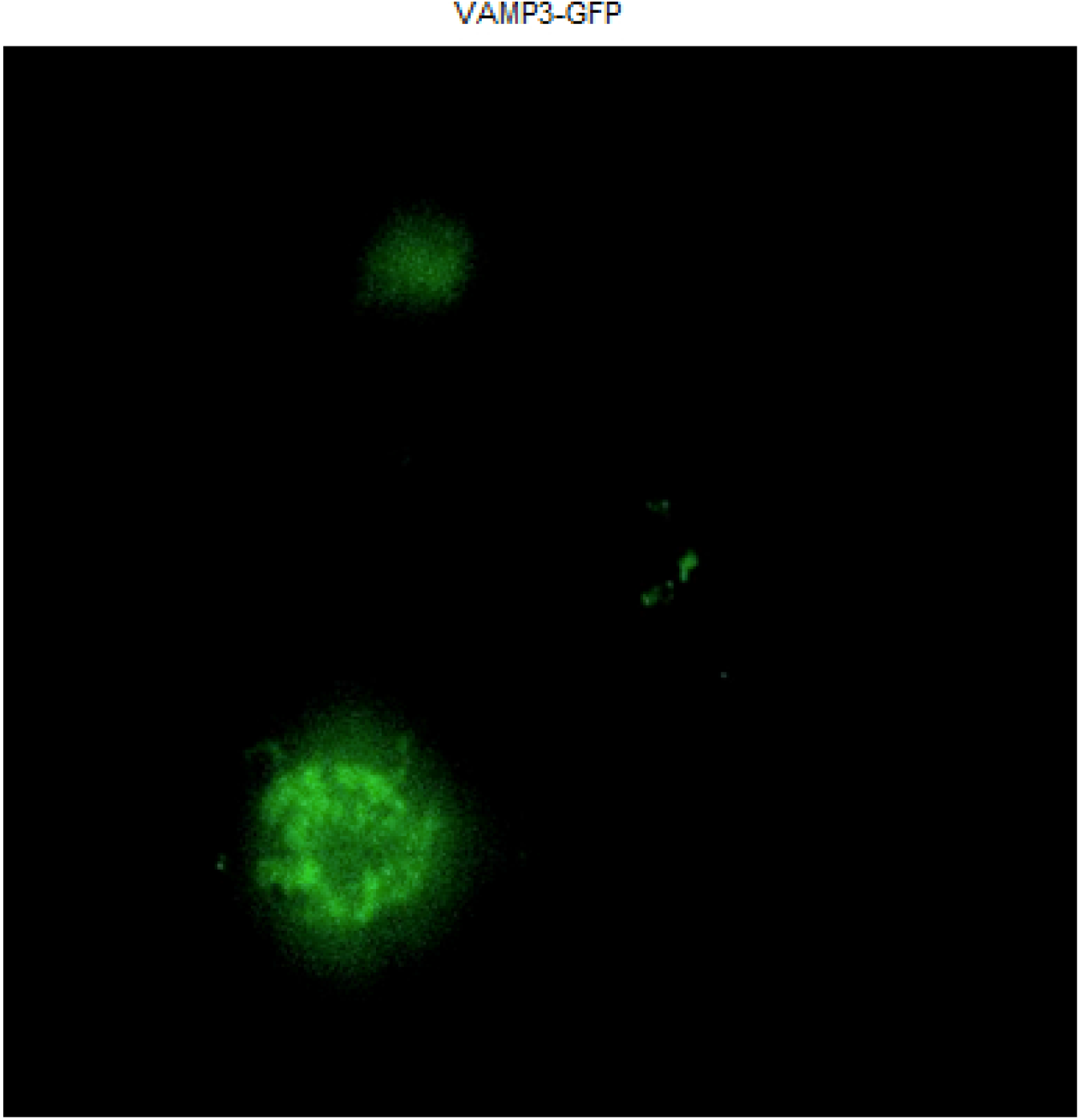
BMDM expressing VAMP3-GFP were followed with time as the cells spread on IgG surfaces, Images were collected every 8 seconds over a 10-minute period with TIRF microscopy

**Video 4:**
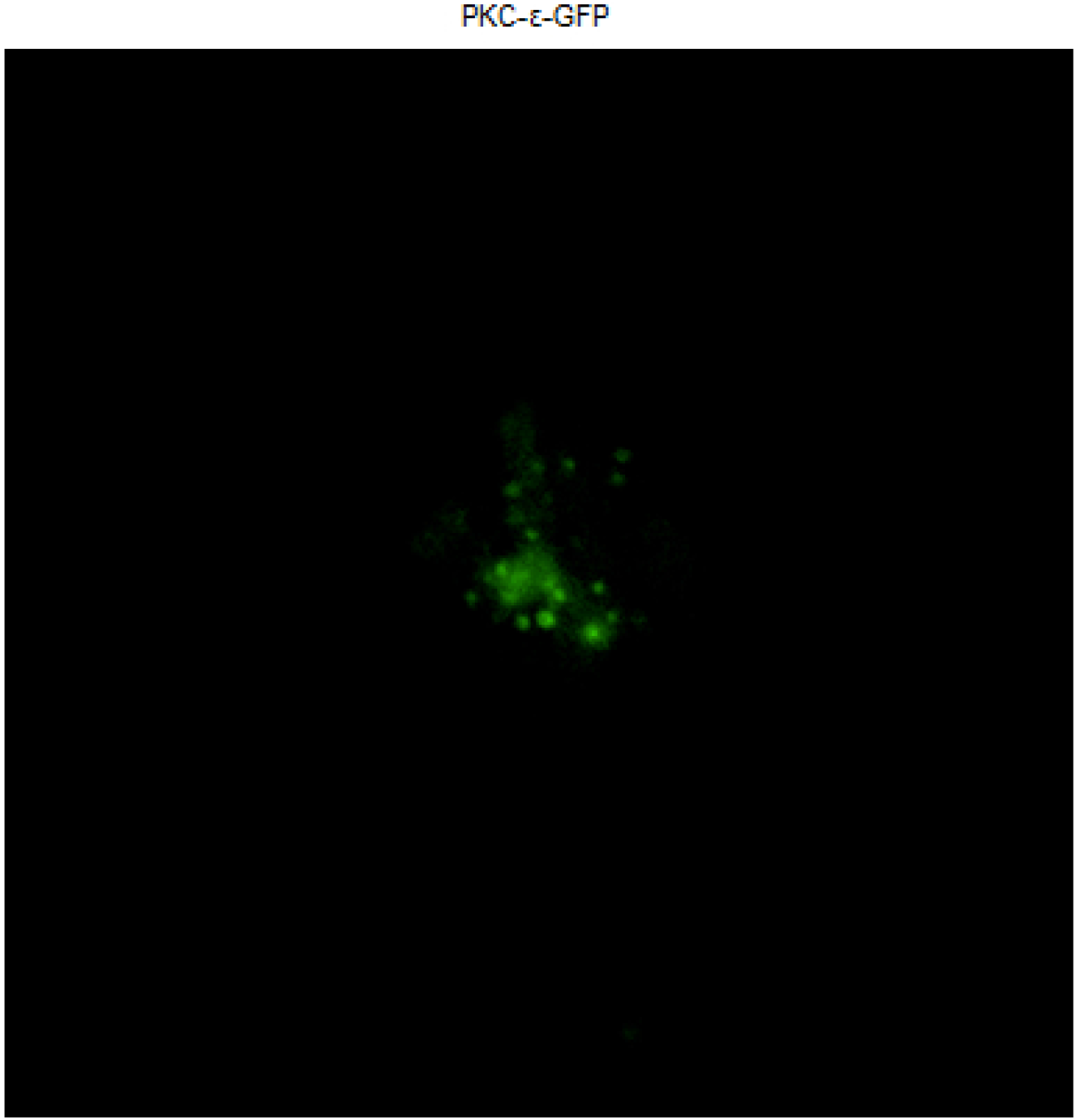
BMDM expressing PKC-ε-GFP were followed with time as the cells spread on IgG surfaces. Images were collected every 8 seconds over a 10-minute period with TIRF microscopy.

**Video 5:**
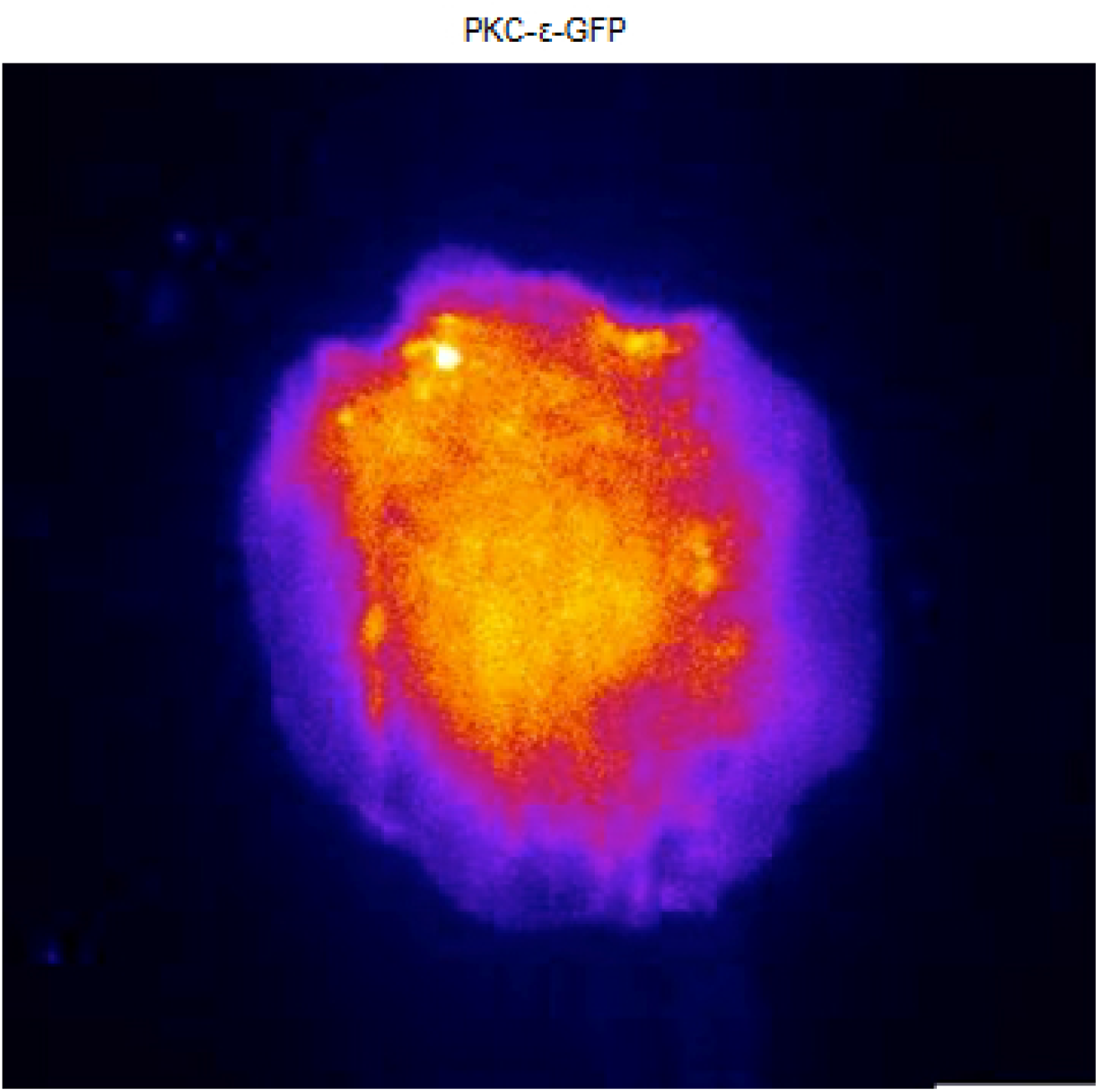
PKC-ε Vesicles Fuse into the Phagosome. BMDM expressing PKC-ε-GFP undergoing frustrated phagocytosis live. Images were collected every 8 seconds over a 10-minute period by TIRF microscopy. Fire LUT was applied for better visualization of fluorescence intensity. Bright puncta representing vesicles are seen entering the phagosome and diffuse into the membrane over time.

**Supplemental Figure 1:**
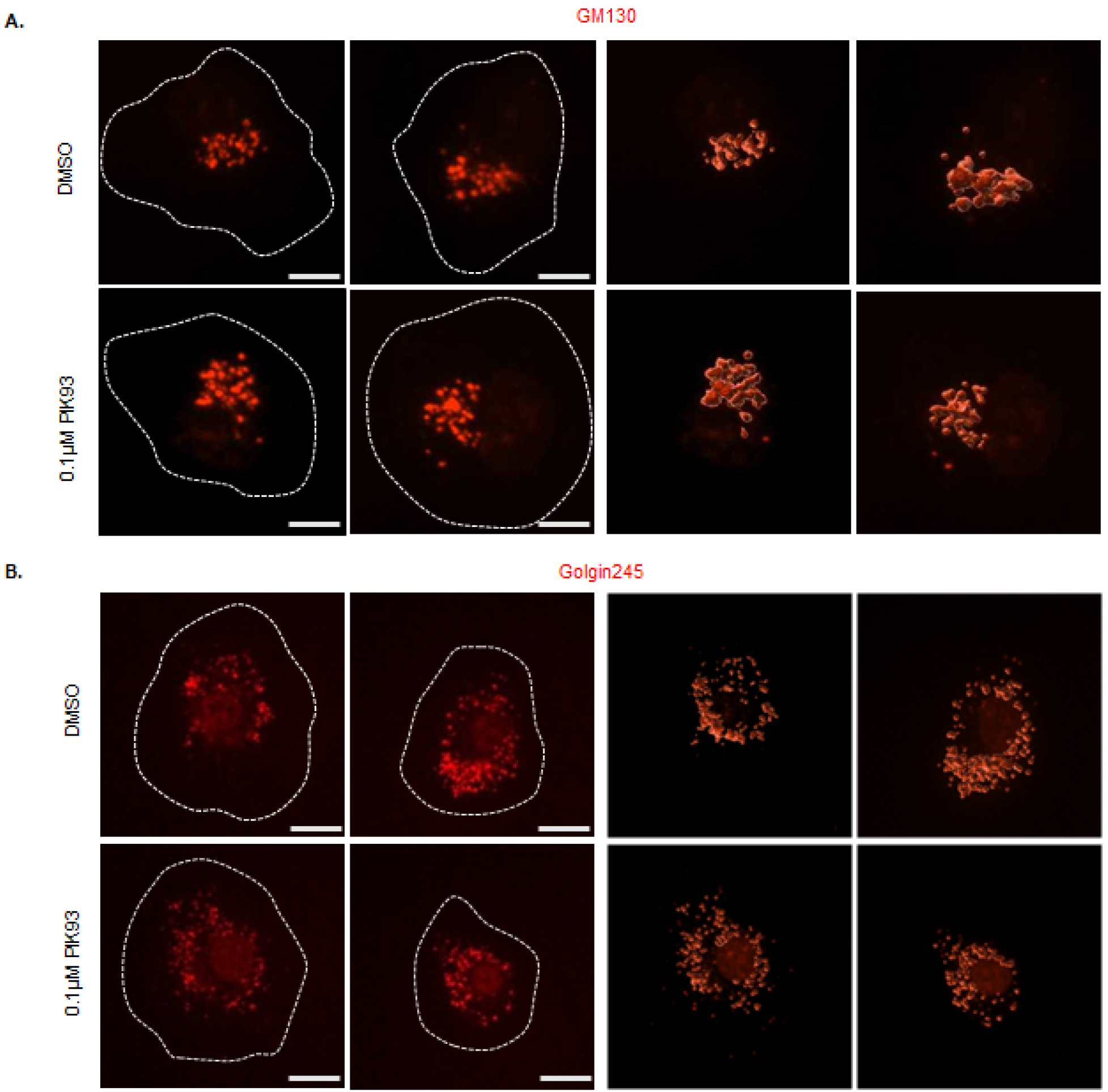
PIK93 Treatment Does Not Alter Golgi Structure. 3D projections of RAW cells treated with DMSO or 100nM PIK93 and stained for (A) cis-medial Golgi marker, GM130, or (B) trans-Golgi marker (Golgin-245). Images were collected with spinning-disc confocal microscopy. Imaris generated 3D renderings visualize overall Golgi structure (right panels). Scale bar = 10 μm. n = 3 (33-36 cells per condition/treatment).

**Supplemental Figure 2:**
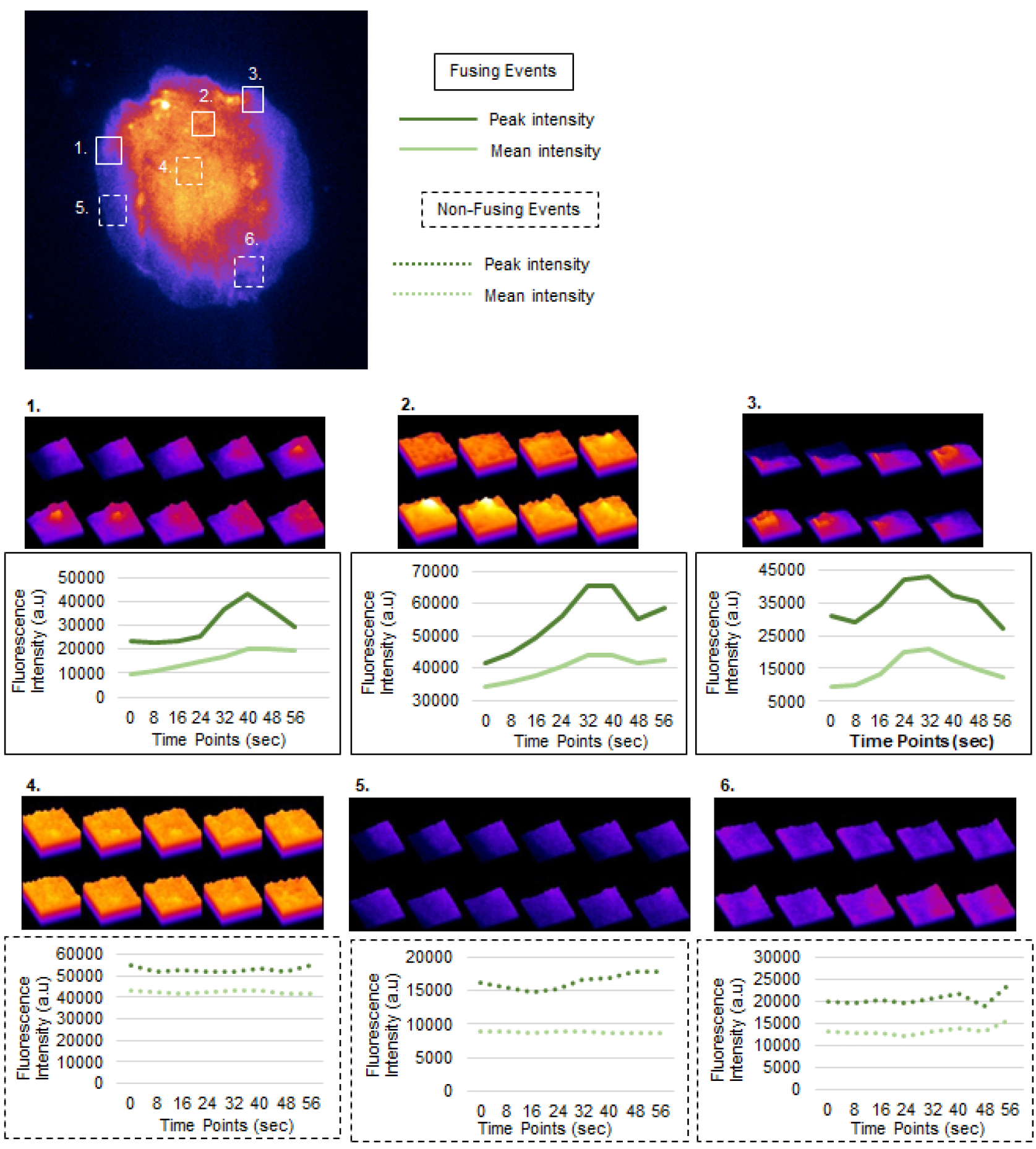
PKC-ε Vesicle Fusion Events. Additional events of fusing (solid boxes) and non-fusing (dotted boxes) from representative cell in Figure 3. 3D surface plots and fluorescent intensity overtime and corresponding measurements of peak and mean fluorescent intensity (graphs) are shown. n = 2 (22 events). Scale bar = 10 μm.

**Supplemental Figure 3:**
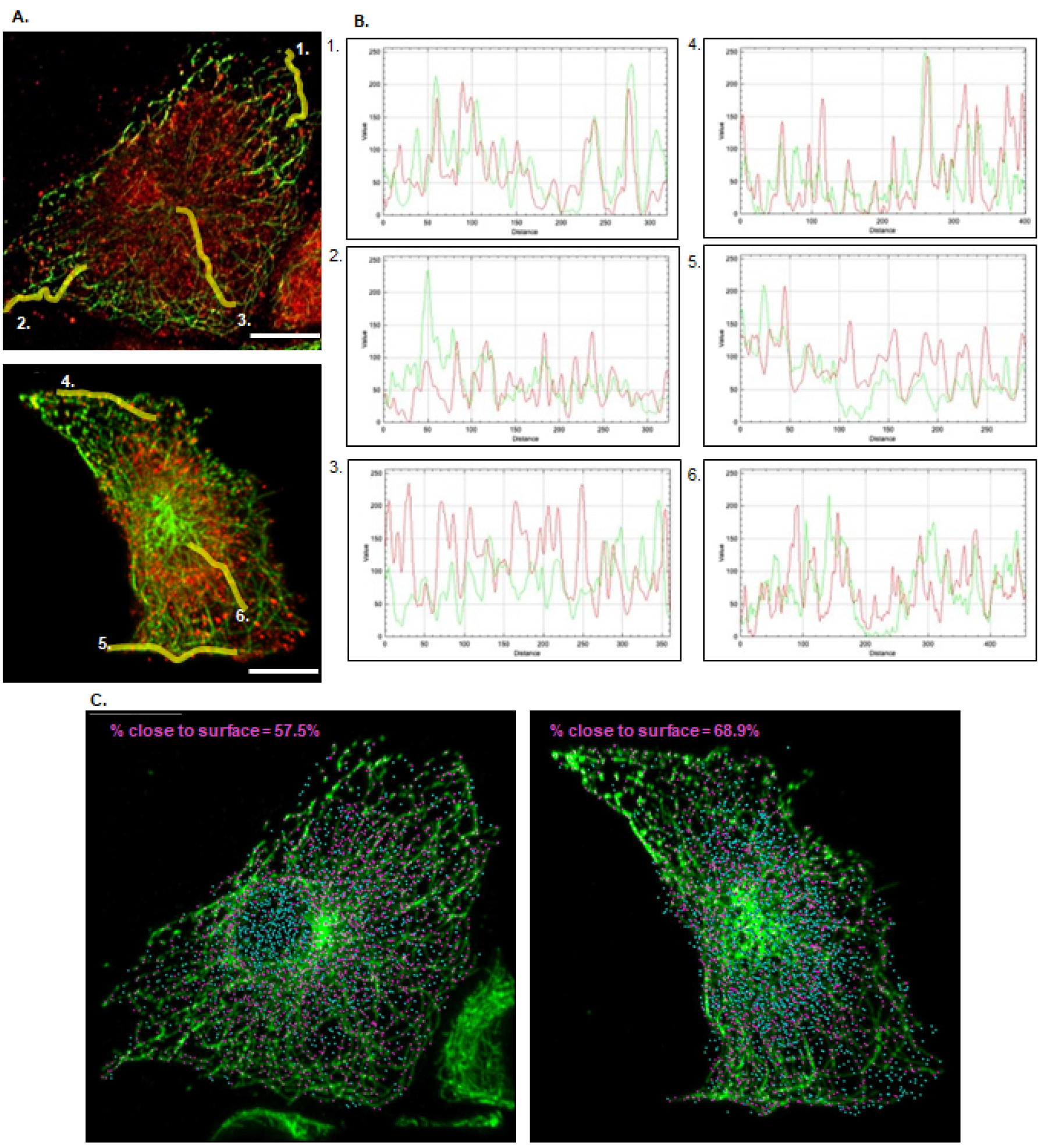
PKC-ε Vesicles Align Along Microtubules. Additional examples of PKC-ε alignment from Figure 5. (A) 3D projections BMDM undergoing frustrated phagocytosis. (B) Line plot analysis of PKC-ε and α-tubulin fluorescence alignment. Line plots presented are of selected microtubules, marked by yellow lines. (D) 3D spot renderings of PKC-ε vesicles aligned (pink spots) along microtubules and PKC-e vesicles not aligned (blue spots) along microtubules. n = 3, 18 cells.

**Supplemental Figure 4:**
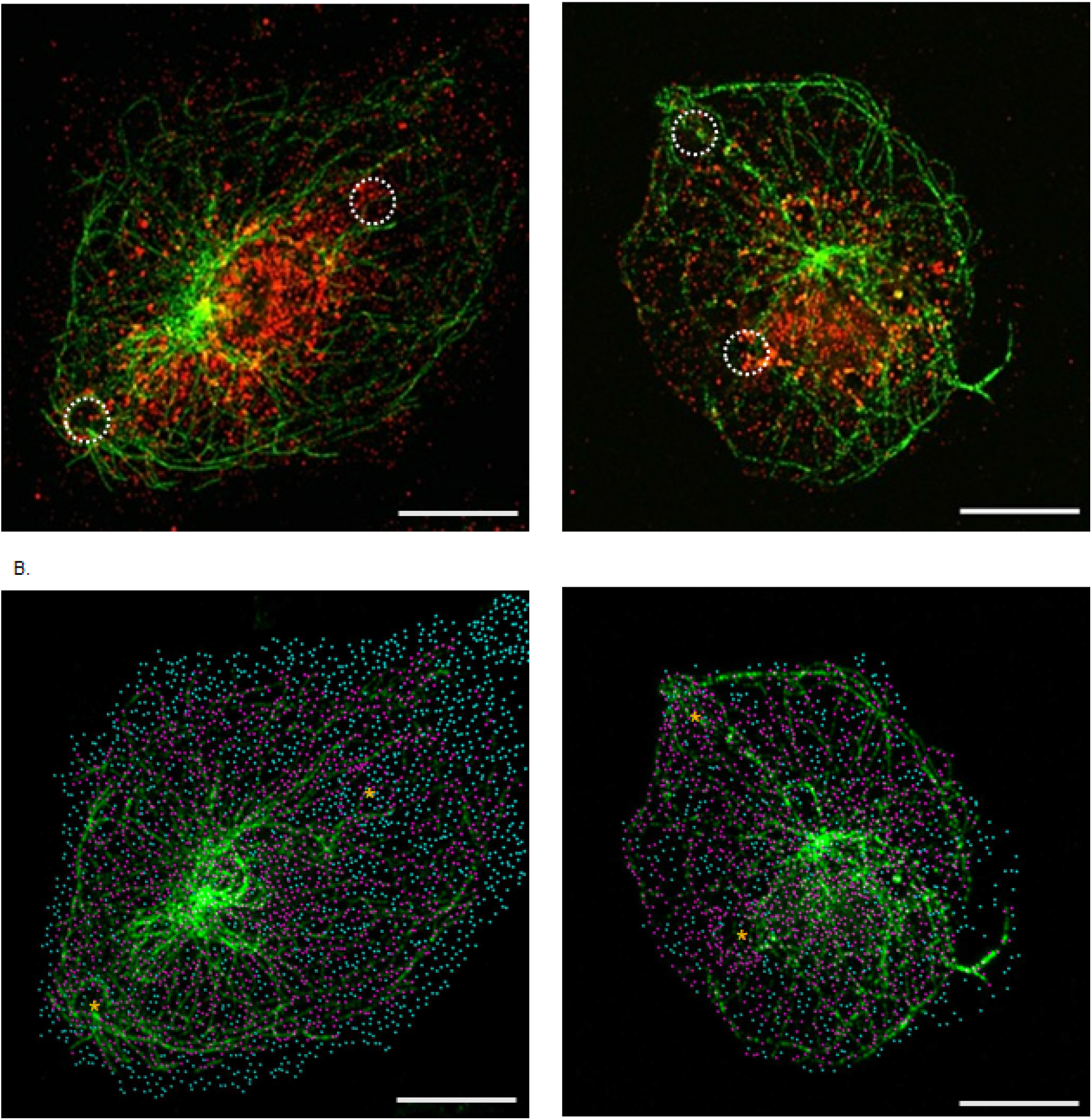
PKC-ε Vesicles Align Along Microtubules Directed towards the Phagosome. Additional examples of PKC-ε alignment on phagosomally directed microtubules from Figure 6. (A) 3D projections of PKC-ε and α-tubulin staining in BMDM phagocytosing 5μm opsonized beads. Dotted circles indicate targets. (B) 3D reconstruction of PKC-ε puncta that are either aligned (pink spots) and non-aligned (blue spots) microtubules. Asterisks indicate targets. n = 3, 57 events.

